# Characterization of the central sulcus pli-de-passage fronto-pariétal moyen in > 1000 human brains

**DOI:** 10.1101/2025.09.11.674690

**Authors:** Anna Marie Muellen, Renate Schweizer

## Abstract

The *pli-de-passage fronto-pariétal moyen* (PPfpm), a deep cerebral fold of the human brain, presents as a common though small elevation at the central sulcus (CS) fundus where it connects the pre- and postcentral gyri at the level of the sensorimotor hand area. Given the PPfpm’s location, single case-reports of its association with the functional sensorimotor hand area, and evidence linking it to the somato-cognitive action network, it holds potential as an anatomical landmark for the sensorimotor region. To characterize the macroscopic morphology of the PPfpm and evaluate its relevance as a reliable and easily detectable anatomical landmark, methods for observer-independent characterization of cortical sulci and structures were adapted and developed to investigate the PPfpm in a large dataset. For 1112 subjects from the Human Connectome Project Young Adult S1200 Release, CS depth profiles were computed from structural magnetic resonance imaging (MRI) data, and an algorithm was developed to automatically extract the PPfpm from these depth profiles. Based on the extraction of two key features approximating the PPfpm at its peak height (PPfpm-I) and its lateral end (PPfpm-II), a principal description of the PPfpm’s position and extent as influenced by hemisphere and sex was conducted. Analyses revealed the PPfpm as a near-universal cerebral fold in the adult human brain, consistently located at mid-height within the CS with a strong association to the CS sulcal pits. Though commonly of a small extent, the PPfpm can be reliably identified in CS depth profiles and in structural MRI data. By providing a systematic, modern macroanatomical characterization of the PPfpm in a large cohort with rigorous quality control, the present study demonstrates the potential of the PPfpm to serve as a robust anatomical landmark for the sensorimotor hand area of the human brain.

## Introduction

First described by Broca (1888), the three *pli-de-passages* of the human central sulcus (CS) are distinct cerebral folds that connect the precentral gyrus of the frontal lobe and the postcentral gyrus of the parietal lobe by traversing the CS - a primary fissure that typically extends continuously from the interhemispheric cleft to the lateral sulcus. Among these *pli-de-passages*, the *pli-de-passage fronto-pariétal supérieur* (PPfps) and the *pli-de-passage fronto-pariétal inférieur* (PPfpi) are prominent, superficial connections that demarcate the superomedial and inferolateral boundaries of the CS. The PPfps is thereby located within the paracentral lobule, while the PPfpi corresponds - in modern terminology - to the subcentral gyrus. While integral to the broader concept of the CS *pli-de-passages*, the PPfps and PPfpi are not the focus of the present study. Rather, this work aims to describe and characterize in detail the third and less well-known structure: the *pli-de-passage fronto-pariétal moyen*.

### Anatomical studies relating to the pli-de-passage fronto-pariétal moyen

Described as a deep fold (“*pli-*”) of small extent (“*toujours très profond*”) passing (“*de-passage*”) from frontal to parietal lobe (“*fronto-pariétal*”) in the middle/at mid-height (“*moyen*”) of the CS (Broca, 1888), the *pli-de-passage fronto-pariétal moyen* (PPfpm) comprises a small and hidden cerebral fold at the fundus of the CS. The PPfpm is shown in a single report (Cunningham, 1890) to arise during intra-uterine brain development where it presents as an “eminence” that initially separates the CS in a shorter upper (superior genu of the CS, ca. 1/3) and a longer lower piece (inferior genu of the CS, ca. 2/3). Parallel to the acceleration of cortical growth around the 5^th^ month of intrauterine brain development and in parallel with the development of the CS between the 20^th^ to 23^rd^ gestational week (Hostalet et al., 2025; Wagner, 1860; White et al., 2010), it is progressively displaced towards the CS fundus while the two CS pieces fuse above it to one continuous fissure (Cunningham, 1890). In the adult brain, the PPfpm persists as a remnant of fetal morphology (Cunningham, 1890), appearing as a small elevation (Cykowski et al., 2008), or a deep annectant gyrus/cerebral fold (“Tiefenwindung”) at the CS fundus at its superomedial third (Eberstaller, 1890; Heschl, 1877; Schweizer et al., 2025).

Due to its concealed location, the PPfpm is less known, and its extent has been characterized in only one large historic dataset from 1877 (Heschl, 1877), a study recently replicated by Schweizer et al. (2025). These investigations confirmed that, while generally of small extent < ⅙ of CS depth, the PPfpm may occasionally remain superficial in the adult brain, thus discontinuing the CS and separating it in its original upper and lower piece. Previously described only in single findings (Alkadhi & Kollias, 2004; Cunningham, 1890; Ecker, 1869; Féré, 1891; McKay et al., 2013; Schweizer et al., 2014; Turner, 1866; Wagner, 1862), two recent studies confirmed the low prevalence of the superficial PPfpm in large cohorts as < 1 % (Mangin et al., 2019; Muellen & Schweizer, 2022; Schweizer et al., 2019, Schweizer et al., 2025). To date, there is no evidence suggesting any pathological significance of this mere anatomical variation.

The anatomy of the CS *pli-de-passages* as cerebral folds traversing the CS was recently confirmed by Skandalakis et al. (2025) through white matter microdissections of 16 human cadaveric brains. Here, the *pli-de-passages* are described as short connections consisting of grey, and primarily white matter. The PPfpm, specifically, was generally located at the level of the middle frontal gyrus, consistent with historical descriptions placing it at the superomedial third of the CS. Studies on the PPfpm in the broader context of CS morphology (Cykowski et al., 2008) in 55 subjects support this finding, confirming that while the PPfpm is morphologically variable, it remains positionally stable and is of generally small extent. Notably, the PPfpm is further implied to be flanked by the sulcal pits of the CS (Im et al., 2010; McKay et al., 2013), developmentally early and spatially stable landmarks within cortical sulci (Hostalet et al., 2025; Lohmann et al., 2008; Régis et al., 2005), suggesting a link between the PPfpm and underlying cortical organization.

Taken together, historical and contemporary studies describe the PPfpm as a common structure within the CS of the human brain, characterized by its small extent and only rare superficial variations, its consistent location at the superomedial third of the CS, and its association with the sulcal pits of the CS. Despite this apparent anatomical consistency, the precise position and extent of the PPfpm has, however, not been systematically characterized in a large cohort. Thus, fundamental questions regarding its precise morphology and the degree of its macroanatomical variability across individuals have not been addressed.

### Association of the *pli-de-passage fronto-pariétal moyen* with functional sensorimotor areas

As described above, the PPfpm is located at the superomedial third of the CS, where it connects the precentral gyrus containing the functional primary motor cortex (M1) and the postcentral gyrus containing the primary somatosensory cortex (S1) (e.g., Penfield & Boldrey, 1937; Penfield & Rasmussen, 1950; Saadon-Grosman et al., 2020) at the level of the functional sensorimotor hand/digit area.

Situated near the “hand knob”, a prominent knob-like folding of the precentral gyrus associated with the functional M1 hand area (Caulo et al., 2007; Rodrigues et al., 2015; Rumeau et al., 1994; Ten Donkelaar et al., 2018; Wagner et al., 2013; Yousry, 1997), the PPfpm may also be associated with sensorimotor hand function. Supporting this, in a rare individual case of a superficial PPfpm discontinuing the CS, the PPfpm was shown to relate to the functional M1 hand area (Alkadhi & Kollias, 2004) during a blood-oxygen-level-dependent functional MRI study. Following finger-to-thumb opposition of the dominant right hand, wrist flexion and extension, and elbow flexion and extension, the superficial PPfpm of this individual was associated with the M1 hand area, and described to separate it from the functional elbow representation. Additionally, in a combined structural and functional MRI study, Germann et al. (2020) reported a close relationship between CS morphology and sensorimotor representations following a set of 14 motor tasks. The PPfpm, herein described as a “submerged gyrus,” was found to lie between two distinct CS segments, separating the sensorimotor representations of the fingers from those of the facial areas. Further evidence of the PPfpm’s association with functional specialization arises from recent work by Gordon et al. (2023) and Skandalakis et al. (2025). Gordon et al. (2023) thereby challenged the classical somatotopic organizational principle of M1 and proposed the “integrate-isolate model”. Within this framework, three effector regions - specific for regional movements - alternate with three inter-effector regions - involved in whole-body action - forming the somato-cognitive action network (SCAN). Given SCAN’s distinct spatial pattern along the precentral gyrus, Skandalakis et al. (2025) proposed that the CS *pli-de-passages* may serve as their neurobiological correlates. The PPfpm thereby corresponds to the middle inter-effector region between the hand and face effector region, highlighting its potential as an anatomical landmark for SCAN and its close spatial relationship with the M1 hand area to which it lies directly lateral. Similarly, Jensen et al. (2023) recently described the “Rolandic motor association” area as a region in the depth of the CS that was observed in 13 subjects to be related to movement coordination. While not explicitly stated and not described with respect to the CS *pli-de-passages*, the location of this area as presented on structural MRI data strongly suggests a link to the PPfpm.

The PPfpm may also relate to the functional S1 hand area: In a clinical Positron-Emission-Topography-study during the presurgical evaluation of 27 tumor and epilepsy patients, Boling and Olivier (2004) showed that peak activation of the functional S1 hand area - elicited through vibrotactile stimulation of the whole hand using a handheld vibrating ball - was located reliably at the PPfpm.

Thus, an increasing body of evidence suggests a close association between the PPfpm and the functional sensorimotor hand/digit area. However, the absence of a detailed anatomical characterization of the PPfpm, along with its limited mention in the existing literature, continues to hinder its broader utilization as an anatomical landmark.

### Aim of the present study

A comprehensive characterization of the PPfpm’s macroanatomy is essential to establish it as a robust anatomical landmark within the central region. Despite its recognized relevance, detailed data on the morphology and interindividual variability of the PPfpm, including the influence of key biological factors, remain sparse. The present study addresses this gap through a systematic, large-scale analysis of the PPfpm’s macroanatomy with rigorous quality control. Consequently, this study aims to enable the reliable identification of the PPfpm, thereby promoting its application as a robust anatomical landmark in both research and clinical practice.

Informed by an established framework for an observer-independent characterization of cortical sulci (Cykowski et al., 2008), a novel method was implemented to extract the PPfpm from structural MRI data in a large cohort of 1112 subjects of the Human Connectome Project Young Adult (HCP-YA) S1200 Release. Following extraction, the macroanatomical extent and position of the PPfpm as influenced by hemispheric location and the subject’s sex were characterized. Lastly, findings were contextualized within the framework of current literature.

## Material and Methods

### Subjects and magnetic resonance imaging data

For 1112 subjects (506 males, 606 females; age in years: 28.80 ± 3.70, range 22 to 37) of the HCP-YA (Van Essen et al., 2013) S1200 Release (Release date: 01 March 2017), unprocessed structural whole brain T1-weighted MRI were downloaded. All data were obtained on a 3 Tesla MRI scanner (Connectome Skyra, Siemens, Erlangen) with a 32-channel head coil and a 3D MPRAGE sequence (TR = 2400 ms; TE = 2.14 ms; TI = 1000 ms; flip angle 8°; iPAT = 2, FOV = 224 x 224 mm²; spatial resolution = 0.7 x 0.7 x 0.7 mm³; TA = 7:40 min).

### Image pre-processing, manual selection and sulcal mesh parametrization of the central sulcus

All data were analyzed with BrainVISA (BrainVISA 4.5.0; Morphologist 2015 Pipeline https://brainvisa.info/). Image pre-processing included manual selection of anterior (AC) and posterior commissure (PC), reorientation to AC-PC-plane and inhomogeneity correction. White and grey matter surface reconstructions were obtained for each hemisphere separately. Within the Morphologist 2015 Pipeline, cortical fold graphs (CFGs) constituting a set of attributed relational graph structures following the medium plane of all sulci (Mangin et al., 2004; Régis et al., 2005) were created for each hemisphere of all subjects. For each hemisphere, the CFGs of the entire hemisphere were visually inspected to identify the CS. An individual CS consists of usually two CFGs (range one to five CFGs), with each single CFG being limited by the anatomical extremities of the sulci - the brain surface and the fundus of the sulci - and constrained by side branches and deep cerebral folds buried within the sulci (Régis et al., 2005). To ensure a correct and complete identification of the CS, all CFGs of each individual CS were selected manually and parametrized with BrainVISA’s Sulcus Parametrization function. The resulting hemispheric datasets consist of a rendered CS mesh (Figure 1 A) and its depth profile (Figure 1 B) for each examined hemisphere. The CS depth profile thereby approximates true CS depth based on the reconstructed CS mesh along its medial to lateral trajectory in a set of 101 discrete data points *i ∈* 1: 101, with the medial end at position *x*_1_ = 0 near the interhemispheric cleft and the lateral end near the lateral sulcus at *x*_101_ = 100. At each of the 101 positions, the depth values *y_i_* are computed as length in mm from the top ridge of the CS mesh to its fundus (Cykowski et al., 2008).

**Figure 1:**
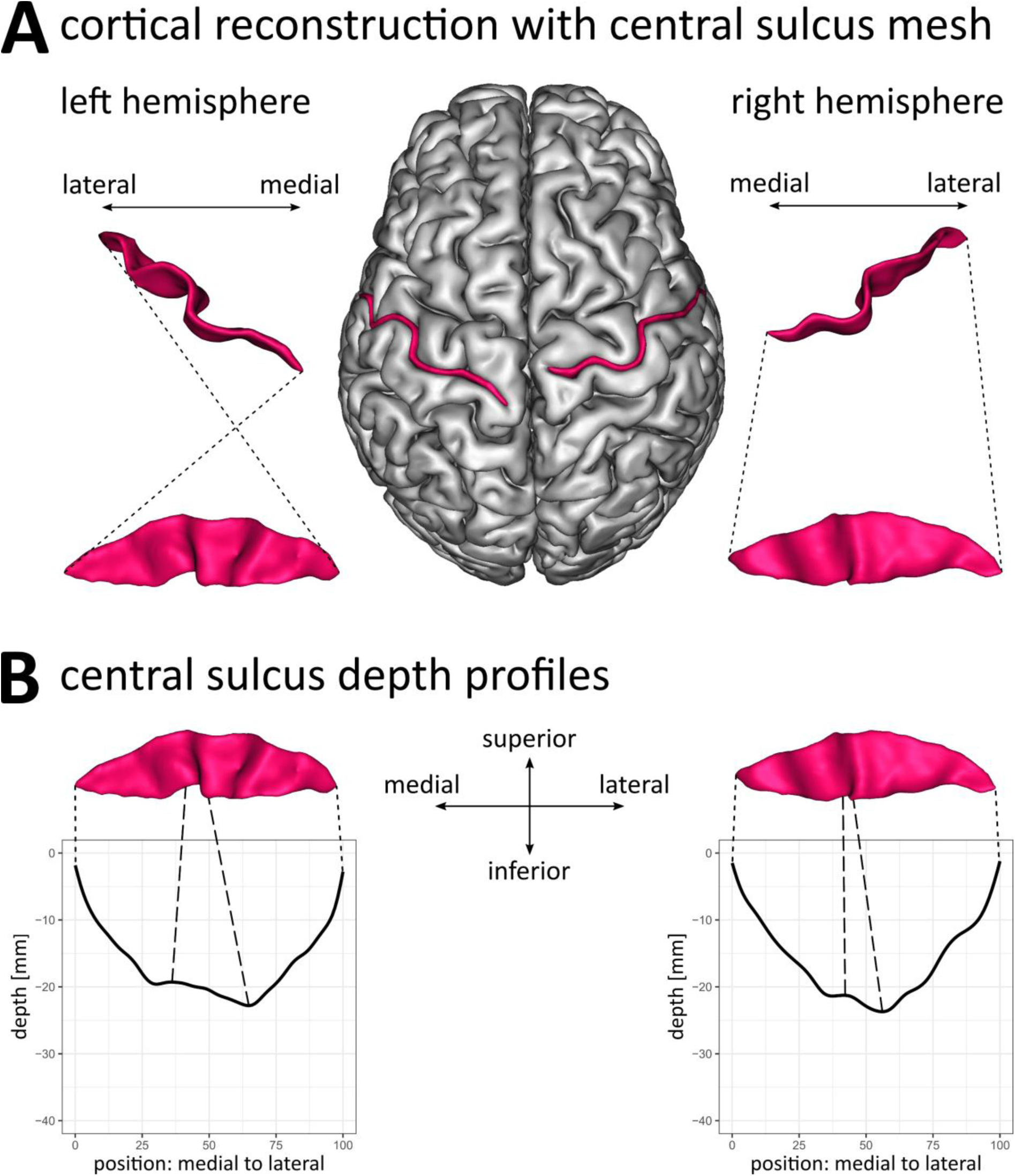
Relation between reconstructed central sulcus meshes and depth profiles. **A** Grey matter surface reconstruction and reconstructed CS mesh (red) of an exemplary subject. **B** Relation between the CS meshes of the left and right hemisphere with their respective depth profiles. Depth values are given at 101 discrete, equidistant position from the medial (position: 0) to the lateral end of the CS (position: 100). Depth values were inverted resulting in a depth profile starting at 0 and going into negative values with increasing depth. Dotted lines mark the equivalent medial and lateral end of reconstructed CS mesh and depth profile. Dashed lines mark the pli de passage fronto-pariétal moyen within the CS mesh and the equivalent point in the depth profile.

### Exclusion of data and quality control

Of the originally obtained 2224 hemispheric datasets from 1112 subjects, 59 hemispheric datasets from 56 subjects were excluded from all further analysis. In 53 subjects, either the left (29 subjects) or the right hemisphere (24 subjects), and for three subjects both the left and right hemispheric datasets were omitted. Reasons for exclusion (multiple can apply to one dataset) included: Data loss during download and computations (5 datasets), incorrect computation of CFGs (29 datasets), incorrect manual selection of CS parts (16 datasets), and anatomical anomalies of the CS which prevented correct computations and selection (30 datasets). Anatomical anomalies included nine cases of a discontinuous CS caused by a superficial PPfpm at brain surface levels (Schweizer et al., 2025). All analyses were performed on the remaining 2165 hemispheric datasets from 1109 subjects (See Table 1, Results).

**Table 1:** Hemispheric datasets after quality control of data.

| groups | subjects |  |  |  |  | hemispheres |  |  |
| --- | --- | --- | --- | --- | --- | --- | --- | --- |
|  | brains (n) | age |  |  |  | datasets (n) |  |  |
| | | M ± SD | SEM | min | max | total | right | left |
| total | 1109 | 28.80 ± 3.70 | 0.11 | 22 | 37 | 2165 | 1085 | 1080 |
| male | 503 | 27.89 ± 3.62 | 0.16 | 22 | 37 | 979 | 491 | 488 |
| female | 606 | 29.56 ± 3.60 | 0.15 | 22 | 36 | 1186 | 594 | 592 |

Abbreviations: M, mean; SD, standard deviation; SEM, standard error of the mean

### Analysis of depth profiles and identification of stable features for the *pli-de-passage fronto-pariétal moyen*

Analysis of 2165 left and right hemispheric depth profiles were performed with the software R (R Core Team (2019) R: A language and environment for statistical computing. R Foundation for Statistical Computing, Vienna, Austria. https://www.R-project.org/.; Version 3.6.1, 05.07.2019 and Version 4.2.2, 31.10.2022, GNU General Public License). For analysis and graphical representation of depth profiles, the depth values *y_i_* were inverted to result in a depth profile starting at 0 and going into negative values with increasing depth. All depth profiles were smoothed (R Core Team (2019). Package stats; function smooth.spline for cubic spline interpolation, generalized cross validation with coefficient of penalty 1, knots defined as all positions *x_i_*; degree of freedom 20).

All extrema of the 2165 depth profiles, i.e., all local and global maxima and minima, were extracted (Package pastecs; function turnpoints; Philippe Grosjean and Frederic Ibanez (2018). pastecs: Package for Analysis of Space-Time Ecological Series. R package version 1.3.21). A two-dimensional kernel density estimation of depth values and positions was computed for all maxima (*n* = 3310), all minima (*n* = 5475), and all global minima (*n* = 2165) to extract clusters of maxima and minima across all depth profiles. Clusters were evaluated on a square grid (100 x 100) defined by each the outermost values (Package MASS; function kde2d; Venables, W. N. & Ripley, B. D. (2002) Modern Applied Statistics with S. Fourth Edition. Springer, New York. R package version 7.3-51.4).

To extract the PPfpm of individual depth profiles, the global minimum at positions > 45 was defined and computed as the depth profile feature “PPfpm-II”, marking the lateral edge of the PPfpm. A second feature “PPfpm-I” was chosen in relation to PPfpm-II as the next medial local maximum to describe the PPfpm at its height. In the infrequent case of a double elevation of the PPfpm, PPfpm-I is thus defined to coherently identify the height of the PPfpm at the directly adjacent maximum to PPfpm-II.

### Quality control of the identified features PPfpm-I and PPfpm-II

Quality was manually assessed for all 2165 hemispheric datasets by one of the authors (AMM) for a thorough and specific validation of the applied method: Assessment of PPfpm-I and PPfpm-II was performed by visual inspection based on a graphical representation of the depth profiles with marked PPfpm-I/-II and compared to the CS meshes. The hemispheric datasets were labelled “correct” when PPfpm-I and PPfpm-II aligned and confirmed clearly with their anatomical equivalent. For all hemispheric datasets where anatomical structures could not be conclusively inferred from depth profiles and CS mesh alone, T1-weighted MR images were included in the assessment. Profiles with at least one feature not aligning with its anatomical equivalent, whether due to anatomical anomalies or failures of the method, where labelled as “incorrect”. Inconclusive hemispheric datasets were marked as “incorrect”. To ensure consistency during the confirmation procedure, previously labelled hemispheric datasets were reassessed in regular intervals. A divergence rate of more than 5 % led to re-evaluation of all respective hemispheric datasets. Interim results on the correctness rates were blinded until the validation procedure was finalized.

### Height and width of the pli-de-passage fronto-pariétal moyen

The absolute height of the PPfpm (PPfpm absolute height) was calculated for all data with both PPfpm-I and PPfpm-II extracted (*n* = 1983, 91.60 %), and defined as their depth difference. The relative height (PPfpm relative height) was defined as the ratio of absolute height to depth at PPfpm-II. Since PPfpm-II coincides in most cases (*n* = 2079, 96.03 %) with the overall deepest point of the CS, this equals a normalization procedure of absolute PPfpm height to maximal CS depth. The width of the PPfpm (PPfpm width) is defined as the absolute difference of positional values at PPfpm-II and PPfpm I. Given the normalization of CS length to positional values 0 to 100, width is thus defined as percentage of total CS length.

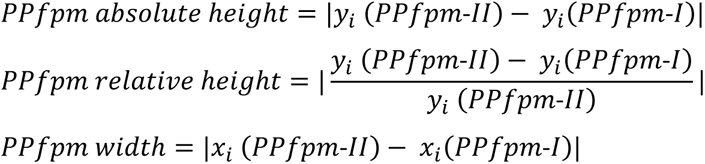

### Statistical analysis of the pli-de-passage fronto-pariétal moyen

All statistical analyses were performed on a subset of 1556 hemispheric datasets from 778 subjects (male: *n* = 374, female: *n* = 404) where both hemispheric datasets exhibit correctly extracted PPfpm-I/-II (Table 3). Descriptive statistics of PPfpm-I/-II values in this subset showed no notable deviation from those of the complete sample.

Three-way mixed-effect-model analysis of variance (ANOVA) (Package ez; function ezANOVA; Michael A. Lawrence (2016). ez: Easy Analysis and Visualization of Factorial Experiments. R package version 4.4.0; type 3 Sum of Squares) was computed to analyze the effect of the “PPfpm feature” (within-subject factor: PPfpm-I, PPfpm-II), “hemisphere” (within-subject factor: right, left), and “sex” (between-subject factor: male, female; given by the HCP-YA as gender, and defined as the stated biological sex of subjects) on position and, separately, on depth at PPfpm-I and PPfpm-II. Normality of distribution for all groups (Shapiro-Wilks, Quantile-Quantile-Plots), and homoscedastic of variance (Levene; cross design on mean for “sex” and “hemisphere” per “feature”) was reviewed. The variable “sex” was not balanced across groups. For all ANOVA effects significant at p < .05, post-hoc two-sided paired t-tests were performed with appropriate Bonferroni correction of the significance threshold p_crit_. Three-way mixed-effect-model ANOVA was conducted to study the effects of “dimension” (within-subject factor: relative PPfpm width, relative PPfpm height), “hemisphere” (within-subject factor: left, right) and “sex” (between-subject factor: male, female) to characterize the PPfpm’s extent further. Supplementary two-way repeated measure ANOVAs were performed on each the relative PPfpm height, the relative PPfpm width, and on the absolute PPfpm height to closer investigate the observed effects of “hemisphere” (within-subject factor: left, right) and “sex” (between-subject factor: male, female). For all analyses, appropriate review of normality of distribution (Shapiro-Wilks, Quantile-Quantile-Plots) and homoscedastic of variance Levene (cross design on mean for sex and hemisphere per type of dimension) was performed. Pearson product-moment correlation coefficient was computed to investigate for potential interdependence and for potential effects of hemispheric origin on relative height and relative width of the PPfpm.

## Results

### The *pli-de-passage fronto-pariétal moyen* can be detected in the depth profiles of the central sulcus

From 1109 subjects (503 male, 606 female; age in years: 28.80 ± 3.70), a total of 2165 CS depth profiles (1080 left hemispheric datasets; 1085 right hemispheric datasets) were surveyed to describe the general pattern of the CS depth trajectory (Table 1), and to identify the PPfpm.

The average outline across all depth profiles comprises a U-shaped trajectory of cortical depth (Figure 2, A). Starting medially at position 0, a moderate decline towards the deepest point at position 61 of −23.04 ± 1.83 mm depth is continued by a comparably steep incline in the lateral third, ending at position 100. Central in the depth profile, a profound but demarked elevation starts in the medial decline and ends in a clearly accentuated drop at the deepest point of the average depth profile, thus discontinuing the otherwise U-shaped depth trajectory. This elevation, the PPfpm, is on average of only limited height. However, a high degree of variations of the PPfpm’s extent and position in the CS depth profile is indicated by the high deviation of depth values at the PPfpm’s maximal height (maximal SD ± 2.49 mm; positions: 45 to 47).

**Figure 2:**
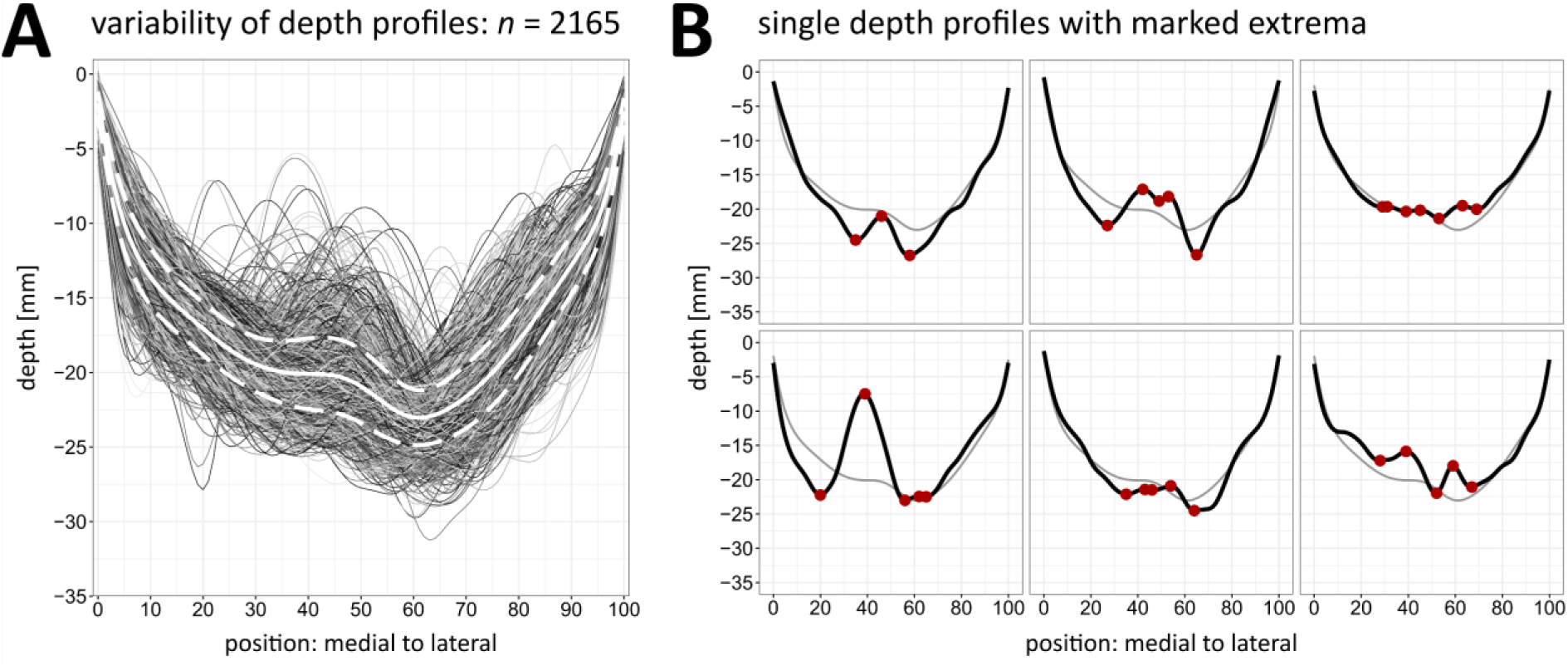
Depth profiles of the central sulcus. Depth [mm] from surface to fundus is given at 101 discrete positions along the CS from medial (x = 0; at the interhemispheric cleft) to lateral (x = 100; at the lateral sulcus). **A** Superimposed depth profiles of 2165 left (n = 1080) and right (n = 1085) hemispheric datasets with average depth (white line) and standard deviation of all depth values (dashed line) computed at each position. **B** Six individual depth profiles: upper row - left hemispheric datasets; bottom row - right hemispheric datasets. Red dots: turn points in individual depth profiles; grey line: average depth profile.

Even though the described depth outline with the centrally located PPfpm can be identified across all 2165 depth profiles, case-by-case inspection stresses the high variability of depth profiles, impeding the identification of common but distinct features for automized PPfpm detection. Individual depth profiles vary in extent and shape of the PPfpm from relatively small (Figure 2 B, upper left) to almost reaching surface levels (Figure 2 B, lower left), including single or double elevations, either pronounced (Figure 2 B, upper middle) or comparatively indistinct (Figure 2 B, lower middle). Depth profiles diverge in overall shape, presenting with a flat trajectory with none or multiple only shallow elevations (Figure 2 B, upper right) or multiple distinct elevations (Figure 2 B, lower right).

### Survey of all depth profile’s extrema indicates a high stability of the global minimum

For an automized extraction of the PPfpm from the depth profiles, the depth profiles’ extrema (*n* = 3310 maxima; *n* = 5475 minima; from *n* = 2165 depth profiles) were identified as stable features, presenting as three prominent clusters along the average depth profile (Figure 3, A). A large, spread-out cluster of maxima is associated with the PPfpm’s maximal height. It is adjoined by and partly overlapping with a distinct medial and a distinct lateral cluster of minima.

**Figure 3:**
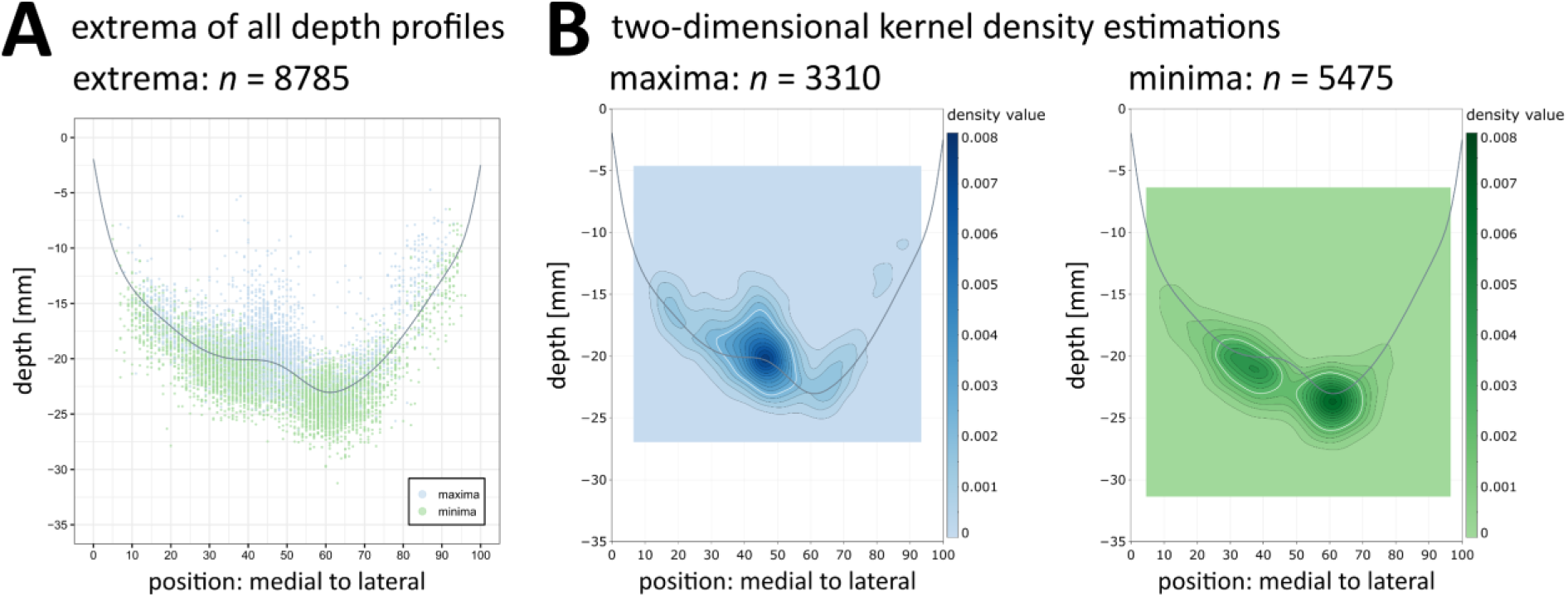
Extrema of all depth profiles (n = 2165). **A** Individual maxima (n = 3310, blue) and minima (n = 5475, green) of 2165 depth profiles. **B** Two-dimensional kernel density estimation of all maxima (left, blue) and all minima (right, green). Two-dimensional kernel density estimation is evaluated on a square grid (100 x 100) as defined by the depth values and positions of each the outer most extrema. Contours of density estimation are given at 0.0005 interval; the white contour line marks the density value 0.0025. Grey line: average depth profile; blue: maxima; green: minima.

To deduce the feasibility of these clusters for automized PPfpm detection and to identify the cluster with the most uniform distribution, two-dimensional kernel density estimations (Figure 3, B) of position vs. depth value were performed for all maxima (Figure 3 B, left) and for all minima (Figure 3 B, right). Density estimation of all maxima revealed a demarked peak incidence distinctly relating to the peak height of the PPfpm in the average depth profile (maximal density value: 0.0072; position 46; depth −20 mm). The estimation, however, implies a high dispersal of position and depth values (density value: 0.0025; positional range: 32 to 55; depth range: 16 mm to −23 mm). Contrastingly, the two-dimensional kernel density estimation of all minima reveals a narrower distribution for both clusters of minima. While the medial cluster is spatially less focused (density value: 0.0025; positional range: 26 to 45; depth range: −19 mm to −23 mm) and comparably attenuated (maximal density value: 0.0046; position: 37; depth: −21 mm), the lateral cluster demonstrates a highly circular density estimation (density value: 0.0025; positional range: 50 to 69; depth range: −21 mm to −26 mm) with a clearly demarked peak density value (maximal density value: 0.0066; position: 61; depth: −24 mm).

Of all three clusters, the lateral cluster of minima is thus suggested to be of a comparably high invariance in both position and depth value. Further evaluation of this cluster against the average depth profile reveals a close correspondence to the deepest point of the average depth profile, thus indicating the lateral cluster to correspond to the deepest points of individual depth profiles. Two-dimensional kernel density estimation of the global minima (Figure 4, A) confirmed a high overlap, revealing a narrow and uniform distribution of global minima with a high peak density value (0.0178; position 61; depth −24 mm). A small divergence from this focalized cluster towards medial positions is caused by 86 global minima at position ≤ 45.

**Figure 4:**
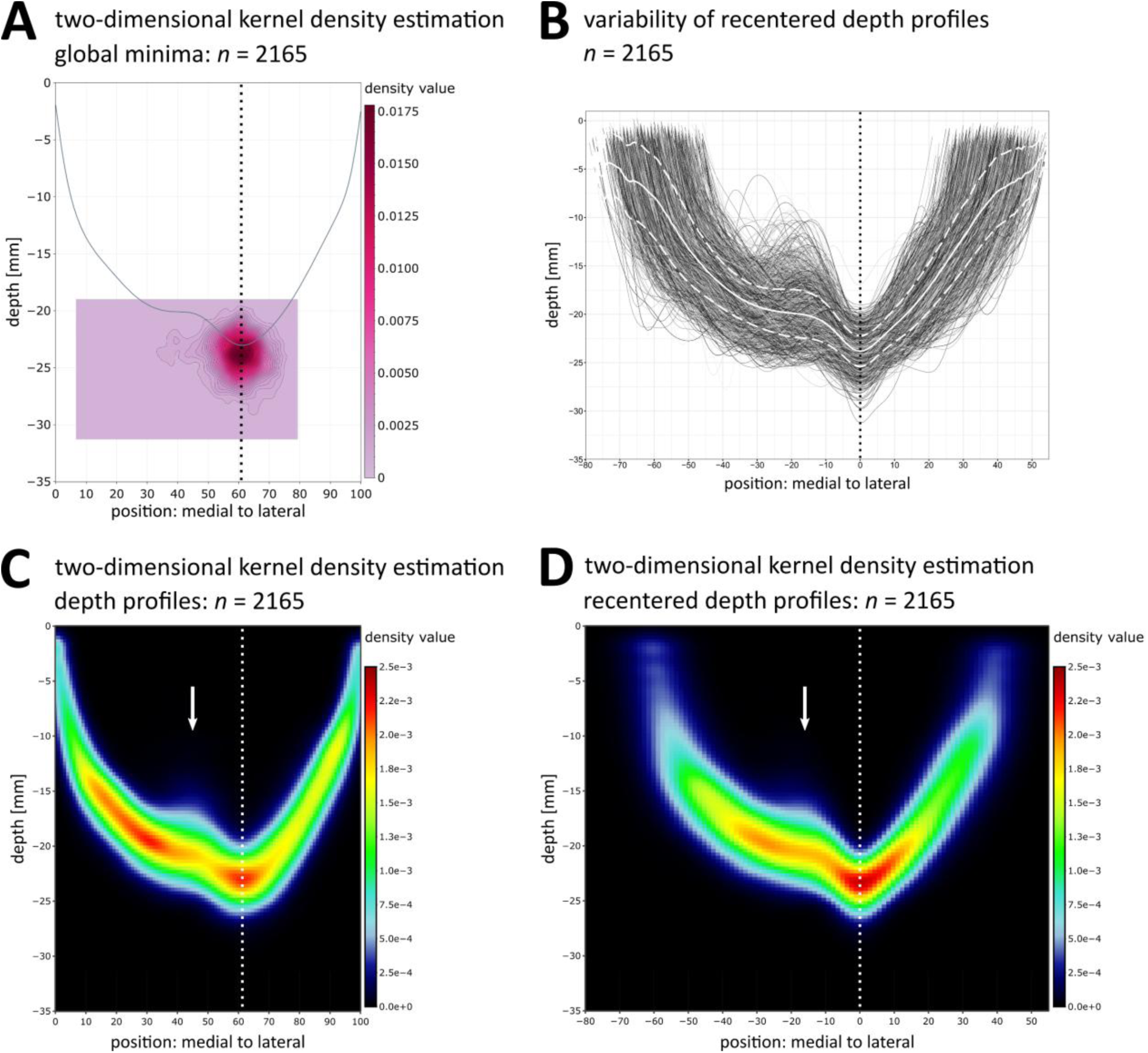
Relation between depth profiles and PPfpm-II (n = 2165). **A** Two-dimensional kernel density estimation of all global minima in depth profiles. Two-dimensional kernel density estimation is evaluated on a square grid (100 x 100) as defined by the depth values and positions of the outermost global minima. Contours of density estimation are given at 0.0005 intervals. Grey line: average depth profile. **B** Superimposed depth profiles after recentering on the respective PPfpm-II (position 0: PPfpm-II; negative values: medial to PPfpm-II; positive values: lateral to PPfpm-II) with average depth (white line) and standard deviation (dashed line) of all depth values computed at each recentered position. PPfpm-II is thereby defined as the deepest local minimum lateral to position 45. **C** Two-dimensional kernel density estimation of all depth profiles before and **D** after recentering on PPfpm-II. Arrow: peak elevation of PPfpm in two-dimensional kernel density estimation across all depth profiles. Dotted line: Average position of PPfpm-II before (A, C: position: 61) and after (B, D: position 0) recentering.

Together, these findings implicate global minima at positions > 45 as highly stable across depth profiles, demarking the PPfpm elevation at its lateral end.

### Identification of stable features of the *pli-de-passage fronto-pariétal moyen*

“PPfpm-II” in depth profiles represents the lateral end of the anatomical pli-de-passage fronto-pariétal moyen

The lateral end of the PPfpm is marked as feature “PPfpm-II” and is defined as the global minima at position > 45 in depth profiles. PPfpm-II is usually located at the deepest point of the CS depth profile. Anatomically, PPfpm-II thus corresponds to the apparently highly stable lateral end of the anatomical PPfpm, usually coinciding with the deepest point of the CS (Figure 5, right).

**Figure 5:**
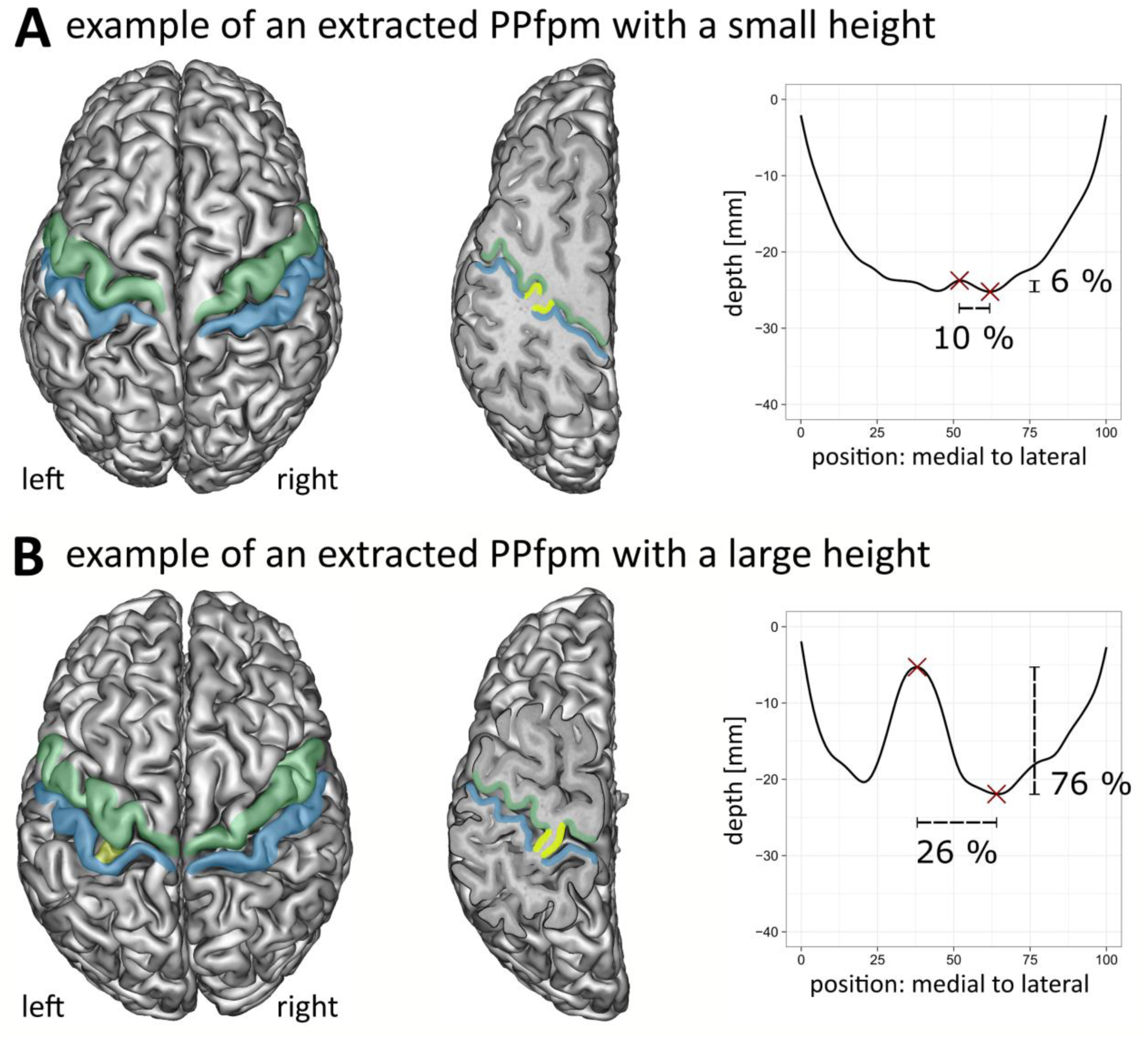
Relation between cortical anatomy and depth profiles. For two exemplary subjects of the HCP-YA, cortical grey matter reconstructions are shown in relation to depth profiles with extracted PPfpm-I / PPfpm-II. Subjects display a PPfpm of **(A)** small and **(B)** large height in the left hemisphere. **Left:** Cortical grey matter reconstructions with marked precentral gyrus (green), postcentral gyrus (blue) and PPfpm (yellow). Note that the PPfpm is not visible in cortical grey matter reconstruction when its height is small, but is apparent when its height is large. **Middle:** Left hemispheric reconstruction with an oblique slicing plane from superior-medial to inferior-lateral in parallel to the CS fundus reveals the PPfpm (yellow) in both subjects. Note that the depth of the oblique slicing plane is different due to the difference in PPfpm height. **Right:** Depth profiles of the left hemisphere with PPfpm-I/-II (red crosses) and with computed relative PPfpm width and height indicated in percent.

Given the observed stability of PPfpm-II, the identification of additional PPfpm-related features in depth profile was anchored on the position of PPfpm-II. The benefit of PPfpm-II as anchor is visualized when individual depth profiles are centered on the PPfpm-II position (Figure 4, B). In these recentered profiles, the PPfpm-elevation is more clearly demarked. While positional and height variability close to the PPfpm peak height (SD ± 2.46 mm; positions: −18 to −19) is retained, the entire PPfpm elevation appears more articulate in comparison to original depth profiles (Figure 2), and - per definition - the average deepest point/lateral end at −23.80 ± 1.60 mm is more accentuated. Two-dimensional kernel density estimations of all depth profiles before (Figure 4, C) and after (Figure 4, D) recentering underline this observation. The density estimation of original depth profiles validates the previously described U-shaped depth trajectory with its focalized deepest point (dotted line) and demonstrates the variability of the PPfpm elevation by a decreased density (arrow). However, after recentering, the PPfpm elevation (arrow) - not the underlying U-shaped depth trajectory - is more homogenous and overall denser.

Thus, extraction of additional PPfpm features is suggested to be more consistent and reliable when defined in relation to PPfpm-II.

“PPfpm-I” in depth profiles represents the peak height of the anatomical pli-de-passage fronto-pariétal moyen

A second depth profile feature, “PPfpm-I”, identifies the peak height of the PPfpm in depth profiles and is extracted as the local maximum medial to PPfpm-II. Since the PPfpm occurs in various shapes, PPfpm-I is defined to consistently identify the peak height directly adjacent to PPfpm-II. Anatomically, PPfpm-I identifies the elevation medially closest to the usually deepest point of the entire CS. PPfpm-I matches the maximal height of the PPfpm in the majority of cases (Figure 5, right).

#### A reliable identification or the medial end of the pli-de-passage fronto-pariétal moyen in depth profiles cannot be attained

Extraction of a third feature demarking the PPfpm towards its medial end was inquired, since the density estimation of minima (Figure 3, C) and the density estimation across all depth profiles (Figure 4, C left) suggested the possibility to define a third feature from depth profile. However, case by case inspection found the medial cluster of minima (Figure 3, C) to include, but not be limited to, features locating to or demarking the medial end of the PPfpm. Figure 2 B provides two examples for this (upper right, lower middle). An accurate match of a third feature to intended anatomical features, i.e., the medial end of the PPfpm, could thus not be guaranteed for an adequate amount of depth profiles in this large data.

#### Quality control of the identified features PPfpm-I and PPfpm-II

The depth profile features PPfpm-I/-II were identified and automatically extracted from 2165 depth profiles to enable an observer-independent characterization of the PPfpm.

Of 2165 analyzed datasets, 1983 depth profiles (91.6 %; left hemispheric datasets: *n* = 1008; right hemispheric datasets: *n* = 975) exhibit both PPfpm-I and PPfpm-II. While PPfpm-II could be identified in all 2165 depth profiles, PPfpm-I is absent in 182 depth profiles (left hemisphere: *n* = 72; right hemisphere: *n* = 110) due to an overall too flat depth profile without any maxima, or exclusion of present maxima as lateral - not medial - to PPfpm-II.

To confirm the matching of both PPfpm-I and PPfpm-II with their anatomical equivalent (Figure 5), individual depth profiles and their extracted features were validated against CS meshes, grey and white matter reconstructions and structural MRI data. Manual quality control revealed an accurate detection of PPfpm-I and PPfpm-II for 2019 datasets (93.3 %; left hemispheric datasets: *n* = 1006; right hemispheric datasets: *n* = 1013), and an erroneous matching of at least one landmark in 143 datasets (6.6 %). For three datasets, a matching could neither be confirmed nor contradicted (0.1 %). Of the 2019 correct depth profiles from 1098 subjects, a subset of 1848 depth profiles (91.53 %) from 1070 subjects showed both PPfpm-I and PPfpm-II extracted correctly (Table 2), enabling a full characterization of the PPfpm.

**Table 2:** Hemispheric datasets after additional quality control of extraction method.

| groups | subjects |  |  |  |  | hemispheres |  |  |
| --- | --- | --- | --- | --- | --- | --- | --- | --- |
|  | brains (n) | age |  |  |  | datasets (n) |  |  |
| | | M ± SD | SEM | min | max | total | right | left |
| total | 1070 | 28.82 ± 3.69 | 0.11 | 22 | 37 | 1848 | 912 | 936 |
| male | 486 | 27.93 ± 3.60 | 0.16 | 22 | 37 | 860 | 424 | 436 |
| female | 584 | 29.56 ± 3.59 | 0.15 | 22 | 36 | 988 | 488 | 500 |

Abbreviations: M, mean; SD, standard deviation; SEM, standard error of the mean

### Characterizing the *pli-de-passage fronto-pariétal moyen* in the depth profiles

Three-way mixed-effect ANOVA statistical analyses were performed on a subset of 778 subjects (*n* = 1556 depth profiles) to examine possible influences on position and depth of PPfpm-I and PPfpm-II and on the derived PPfpm extent (width, height) by sex (male, female) and hemisphere (left, right). For these analyses, subjects were considered only if both PPfpm-I and PPfpm-II were extracted correctly for both hemispheres (Table 3).

**Table 3:** Hemispheric datasets for statistical analysis. Both left and right hemispheric datasets exhibit a correctly extracted PPfpm-I and PPfpm-II.

| groups | subjects |  |  |  |  | hemispheres |  |  |
| --- | --- | --- | --- | --- | --- | --- | --- | --- |
|  | brains (n) | age |  |  |  | datasets (n) |  |  |
| | | M ± SD | SEM | min | max | total | right | left |
| total | 778 | 28.85 ± 3.70 | 0.13 | 22 | 37 | 1556 | 778 | 778 |
| male | 374 | 28.09 ± 3.61 | 0.19 | 22 | 37 | 748 | 374 | 374 |
| female | 404 | 29.54 ± 3.65 | 0.18 | 22 | 36 | 808 | 404 | 404 |

Abbreviations: M, mean; SD, standard deviation; SEM, standard error of the mean

#### Position and depth of the depth profile features PPfpm-I/-II

PPfpm-I is on average located in the medial half of the CS (position 45.77 ± 6.12) with an average depth of −19.25 ± 2.74 mm. PPfpm-II is located at the lateral third of the CS (position 60.79 ± 5.12) with an average depth of −23.78 ± 1.60 mm. PPfpm-II closely correlates with the deepest point of the CS at position 59.67 ± 7.32 and with an average depth of −23.82 ± 1.59 mm (Table 4).

**Figure 6:**
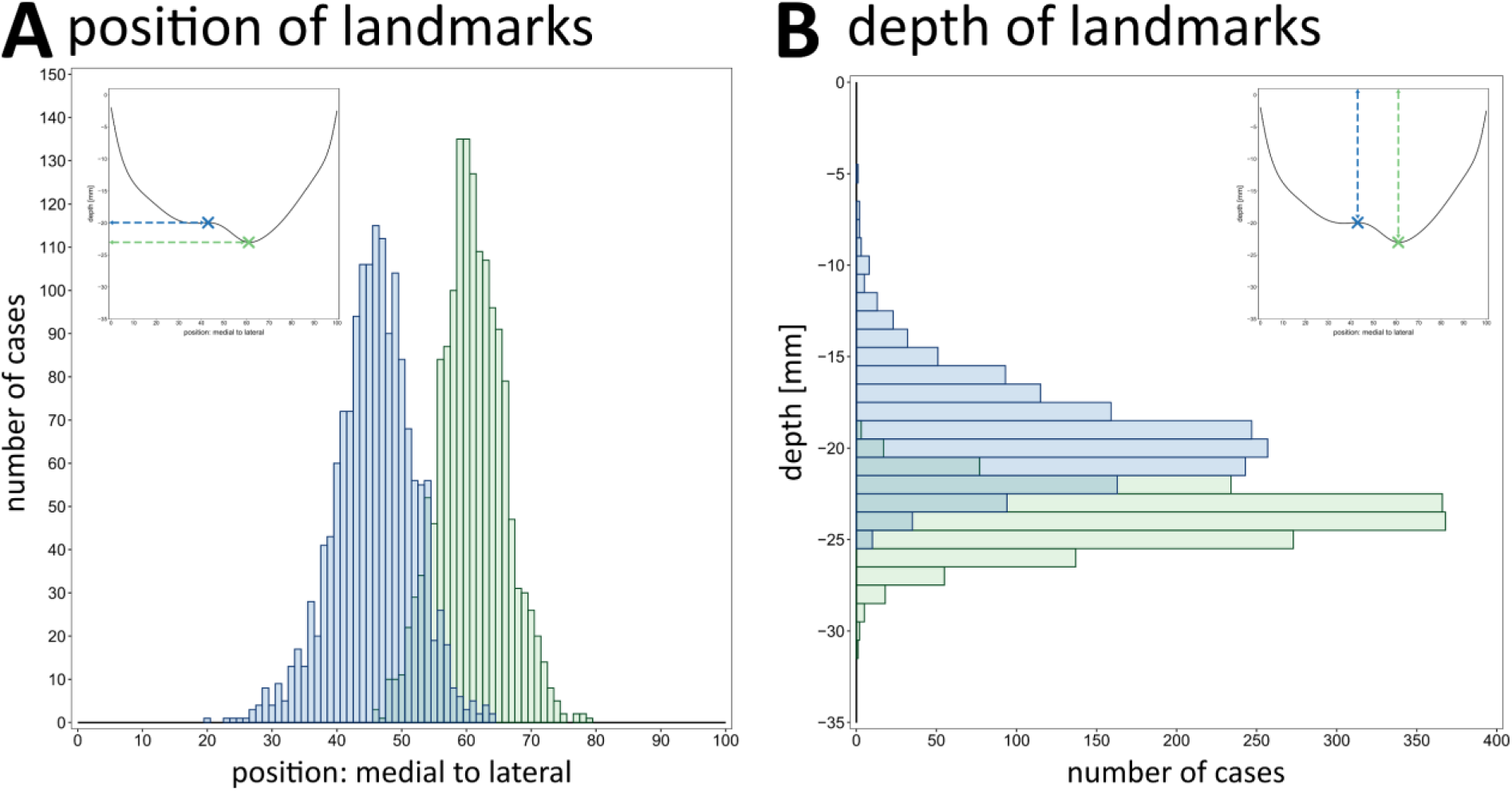
Position and depth values of PPfpm-I/-II. Position **(A)** and depth [mm] **(B)** of PPfpm-I (blue) and PPfpm-II (green) for 778 subjects (n = 1556 depth profiles) where both are extracted correctly in both hemispheres. Inserts of average depth profile with marked PPfpm-I/-II are shown in the upper corners for comparison.

**Table 4:** Descriptive statistics of depth profiles (*n* = 1556 datasets).

|  | position |  |  | depth [mm] |  |  |
| --- | --- | --- | --- | --- | --- | --- |
|  | PPfpm-I | PPfpm-II | deepest point | PPfpm-I | PPfpm-II | deepest point |
| M ± SD | 45.77 ± 6.12 | 60.79 ± 5.12 | 59.67 ± 7.32 | -19.25 ± 2.74 | -23.78 ± 1.60 | -23.82 ± 1.59 |
| SEM | 0.16 | 0.13 | 0.19 | 0.07 | 0.04 | 0.04 |
| min | 20 | 46 | 7 | -5.30 | -19.04 | -19.04 |
| max | 64 | 79 | 79 | -25.17 | -31.24 | -31.24 |

Abbreviations: M, mean; SD, standard deviation; SEM, standard error of the mean

#### Position of PPfpm-I/-II

Analyses on the features’ position (Table 5) confirms PPfpm-I - per definition - significantly more medial to PPfpm-II. Both PPfpm-I and PPfpm-II are more lateral in right hemispheres (Figure 7, A) with the effect more apparent on PPfpm-I (PPfpm-I: Δ = 2.21 positions; PPfpm-II: Δ = 1.38 positions; Bonferroni correction: p_crit_ < .0125; Table 6). In contrast, PPfpm-II demonstrates a higher positional stability across hemispheres and subjects as indicated by the smaller effect size and the overall narrower distribution.

**Figure 7:**
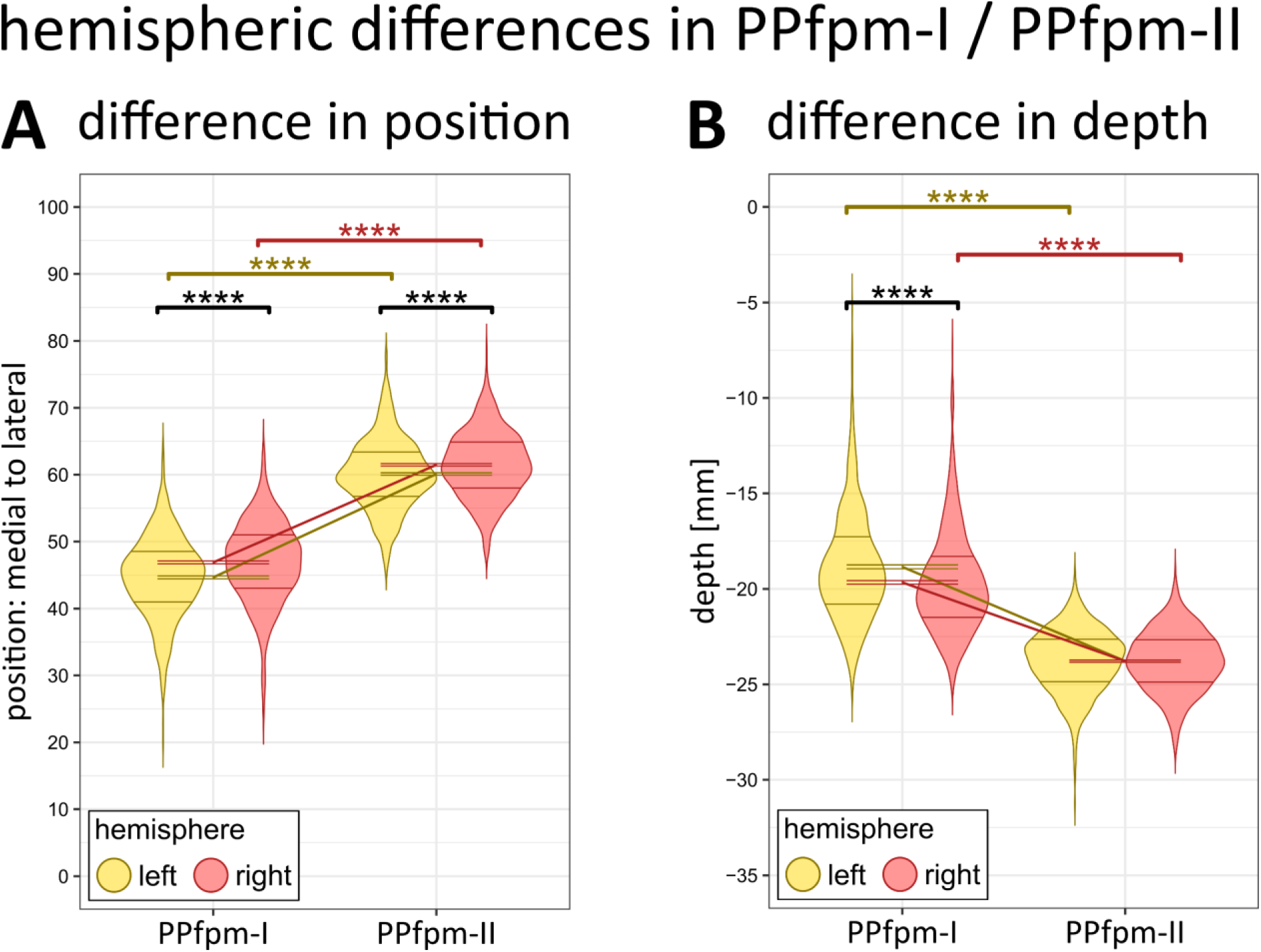
Hemispheric differences (left, right) in PPfpm-I and PPfpm-II, each in their position (A) and in their depth (B). Distribution of individual values is shown as violin plot (Gaussian kernel) with .25 and .75 quartile indicated by horizontal lines; M ± SEM are given by hemisphere for PPfpm-I/-II. Significance of effects: Post-hoc paired t-test (Bonferroni correction: p_crit_<.0125) indicate a significant effect of hemisphere and feature on positional and depth values: **** p<.000025, *** p<.00025, ** p<.0025, * p<.0125, ns not significant. Yellow: left hemisphere; red: right hemisphere.

**Table 5:**
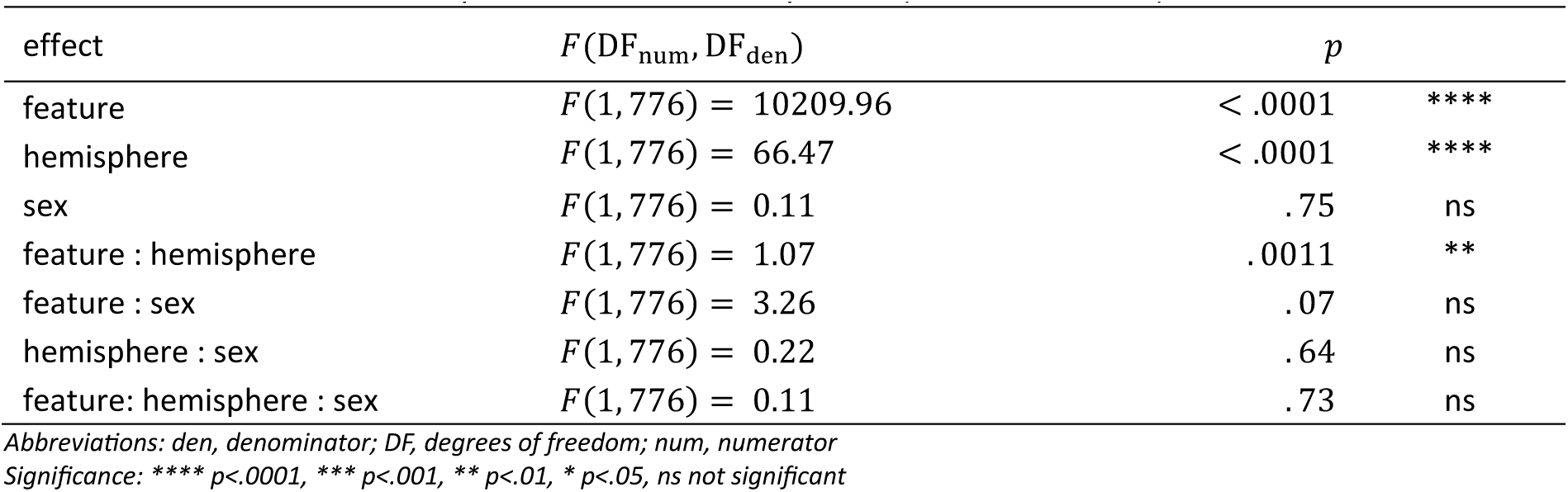
Mixed-effect ANOVA on position of features PPfpm-I/-II (*n* = 1556 datasets).

**Table 6:** Bonferroni corrected post-hoc testing of position of features PPfpm-I/-II per hemisphere (*n* = 1556 datasets).

| condition | comparison | position |  |  |  | paired two-sided t-test |  |  | 95 % confidence interval |  |
| --- | --- | --- | --- | --- | --- | --- | --- | --- | --- | --- |
| | | M | SD | SEM | $\Delta$ | t | $p$ | | lower | upper |
| PPfpm-I | left | 44.67 | 6.03 | 0.22 | -2.21 | -7.86 | $< .0001$ | **** | -2.76 | -1.66 |
|  | right | 46.88 | 6.02 | 0.22 |  |  |  |  |  |  |
| PPfpm-II | left | 60.10 | 5.22 | 0.19 | -1.38 | -6.15 | $< .0001$ | **** | -1.81 | -0.94 |
|  | right | 61.48 | 4.93 | 0.18 |  |  |  |  |  |  |
| left | PPfpm-I | 44.67 | 6.03 | 0.22 | -15.44 | -79.97 | $< .0001$ | **** | -14.99 | -14.21 |
|  | PPfpm-II | 60.10 | 5.22 | 0.19 |  |  |  |  |  |  |
| right | PPfpm-I | 46.88 | 6.02 | 0.22 | -14.60 | -73.81 | $< .0001$ | **** | -15.81 | -15.06 |
|  | PPfpm-II | 61.48 | 4.93 | 0.17 |  |  |  |  |  |  |

Abbreviations: $\Delta$ , estimation of difference of means between comparisons; M, mean; SD, standard deviation; SEM, standard error of the mean Bonferroni adjusted significance: \*\*\*\* $p < .000025$ , \*\*\* $p < .00025$ , \*\* $p < .0025$ , \* $p < .0125$ , ns not significant

#### Depth at PPfpm-I/-II

Analyses of the depth value (Table 7) implicates PPfpm-II - per extraction method - significantly deeper than PPfpm-I. Depth values at PPfpm-II are indicated as stable across hemisphere (Figure 7, B), while depth at PPfpm-I is significantly deeper in right hemispheres (PPfpm-I: Δ = 0.82 mm; Bonferroni correction p_crit_ < .0125; Table 8).

**Table 7:** Mixed-effect ANOVA on depth of features PPfpm-I/-II (*n* = 1556 datasets).

| effect | $F(DF_{\text{num}}, DF_{\text{den}})$ | $p$ | |
| --- | --- | --- | --- |
| feature | $F(1, 776) = 3765.22$ | $< .0001$ | **** |
| hemisphere | $F(1, 776) = 45.06$ | $< .0001$ | **** |
| sex | $F(1, 776) = 81.54$ | $< .0001$ | **** |
| feature : hemisphere | $F(1, 776) = 4.40$ | $< .0001$ | **** |
| feature : sex | $F(1, 776) = 1.00$ | .32 | ns |
| hemisphere : sex | $F(1, 776) < .0001$ | 1.00 | ns |
| feature : hemisphere : sex | $F(1, 776) = 1.08$ | .30 | ns |

Abbreviations: den, denominator; DF, degrees of freedom; num, numerator Significance: \*\*\*\* $p < .0001$ , \*\*\* $p < .001$ , \*\* $p < .01$ , \* $p < .05$ , ns not significant

**Table 8:** Bonferroni corrected post-hoc testing of depth at features PPfpm-I/-II per hemisphere (*n* = 1556 datasets).

| condition | comparison | depth value [mm] |  |  |  | paired two-sided t-test |  |  | 95 % confidence interval |  |
| --- | --- | --- | --- | --- | --- | --- | --- | --- | --- | --- |
| | | M | SD | SEM | $\Delta$ | t | $p$ | | lower | upper |
| PPfpm-I | left | -18.84 | 2.80 | 0.10 | 0.82 | 7.43 | $< .0001$ | **** | 0.60 | 1.03 |
|  | right | -19.66 | 2.62 | 0.09 |  |  |  |  |  |  |
| PPfpm-II | left | -23.78 | 1.64 | 0.06 | 0.00 | -0.04 | .97 | ns | -0.1 | 0.1 |
|  | right | -23.78 | 1.57 | 0.06 |  |  |  |  |  |  |
| left | PPfpm-I | -18.84 | 2.80 | 0.10 | 4.94 | 49.50 | $< .0001$ | **** | 4.74 | 5.14 |
|  | PPfpm-II | -23.78 | 1.64 | 0.06 |  |  |  |  |  |  |
| right | PPfpm-I | -19.66 | 2.62 | 0.09 | 4.12 | 45.16 | $< .0001$ | **** | 3.94 | 4.30 |
|  | PPfpm-II | -23.78 | 1.57 | 0.06 |  |  |  |  |  |  |

Abbreviations: $\Delta$ , estimation of difference of means between comparisons; M, mean; SD, standard deviation; SEM, standard error of the mean Bonferroni adjusted significance: \*\*\*\* $p < .000025$ , \*\*\* $p < .00025$ , \*\* $p < .0025$ , \* $p < .0125$ , ns not significant

Taken together with the statistical analysis of positions at PPfpm-I/PPfpm-II, right hemispheres are implicated to exhibit flatter depth profiles with an - on average - deeper and more lateral peak height.

#### Extent of the pli-de-passage fronto-pariétal moyen in depth profiles

The extent of the PPfpm, i.e., the width and height, is derived from the depth profile features PPfpm-I/-II (Figure 5). The PPfpm width as the total positional difference of PPfpm-II and PPfpm-I is given as a relative width of total CS length (15.02 ± 5.47 %). As such, relative PPfpm width is defined between the PPfpm’s lateral end (PPfpm-II) and peak height (PPfpm-I), and not extended to the more intuitive - although unreliable - medial end of the PPfpm. The PPfpm height is the absolute difference in depth between PPfpm-II and PPfpm-I (4.53 ± 2.70 mm). Single case normalization of the absolute PPfpm height to depth at PPfpm-II gives a relative PPfpm height of 18.94 ± 11.07 % (Table 9).

**Figure 8:**
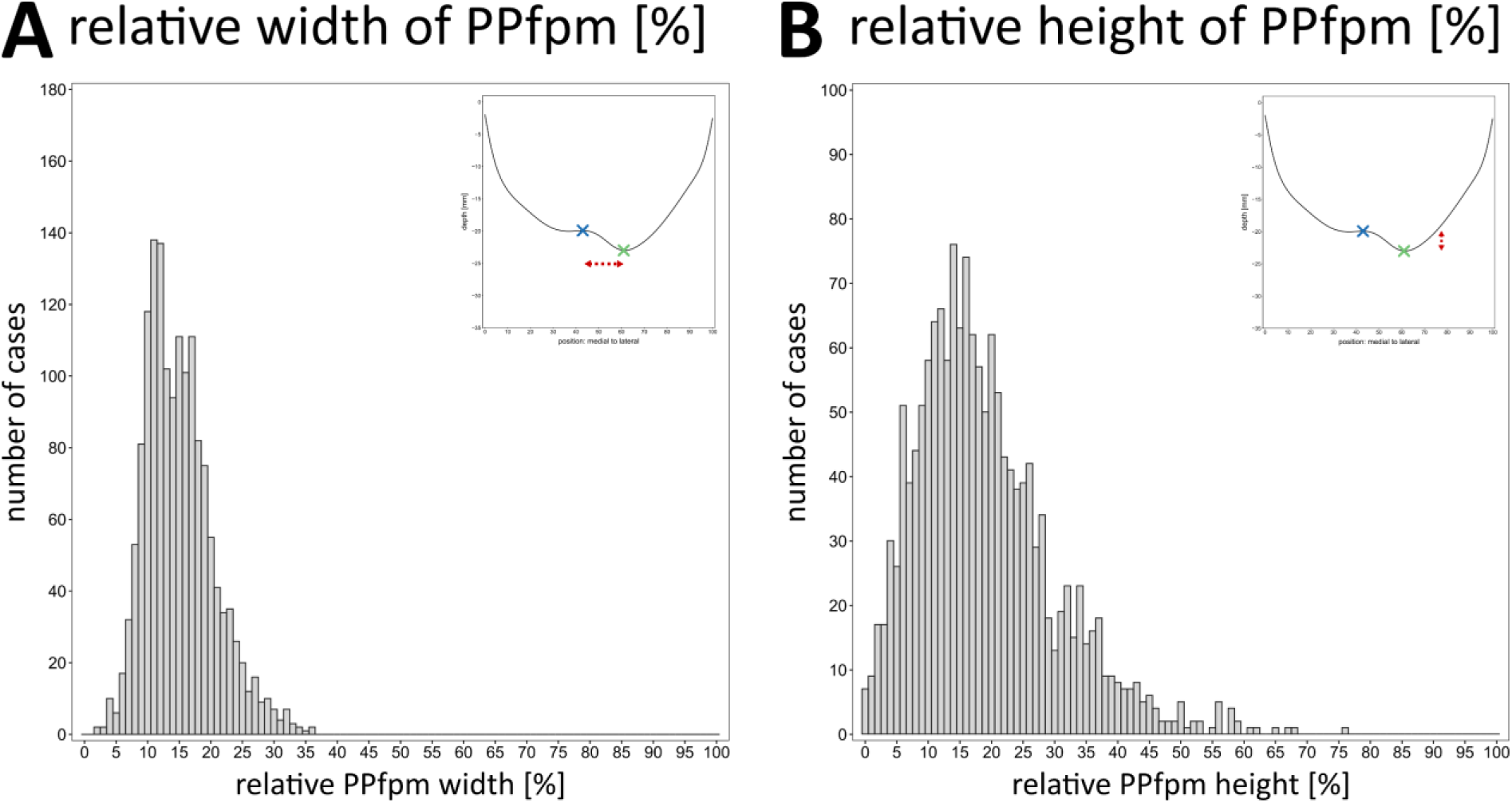
Extent of the PPfpm. Width **(A)** and height **(B)** of the PPfpm are given in relation to total CS length and depth at PPfpm-II. Upper right corner shows the average depth profile for comparison with marked PPfpm-I (blue) and PPfpm-II (green), and the observed dimension (red arrow-line).

**Table 9:** Descriptive statistics of PPfpm extent (*n* = 1556 datasets).

|  | relative width [%] | absolute height [mm] | relative height [%] |
| --- | --- | --- | --- |
| M ± SD | 15.02 ± 5.47 | 4.53 ± 2.70 | 18.94 ± 11.07 |
| SEM | 0.14 | 0.07 | 0.28 |
| min | 2.00 | 0.02 | 0.06 |
| max | 36.00 | 16.64 | 75.85 |

Abbreviations: M, mean; SD, standard deviation; SEM, standard error of the mean

#### Relative width and relative height of the pli-de-passage fronto-pariétal moyen correlate strongly

Investigating possible interdependencies of relative PPfpm height and width, Pearson correlation coefficient was found strong at *R* = .63, *p* (two-tailed) < .0001 across all datasets, confirming a higher PPfpm to also be broader from peak height to lateral end.

Considering the higher stability of the PPfpm’s lateral end (PPfpm-II), this indicates the PPfpm’s peak height (PPfpm-I) to be shifted to medial positions the higher it is. Separate analyses by hemisphere (Figure 9, A) revealed no relevant difference in correlation, suggesting the effect to be universal across hemispheres.

**Figure 9:**
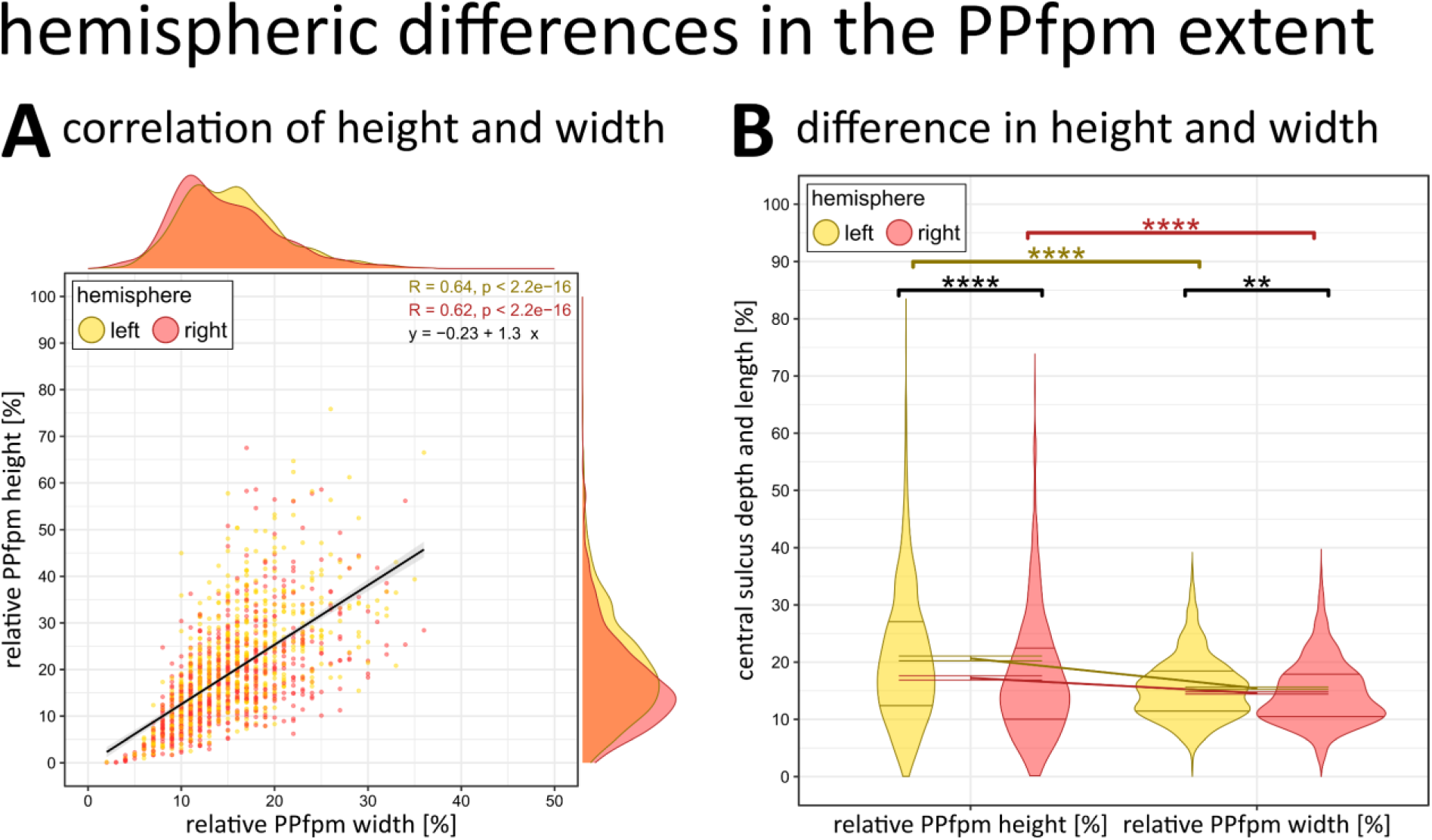
Hemispheric differences (left, right) in the PPfpm extent. **(A)** Correlation of relative PPfpm height [%] and relative PPfpm width [%] for 778 subjects (n = 1556 datasets). Note that width is defined as positional difference between PPfpm-II and PPfpm-I, and thus equals a width from the anatomical PPfpm’s lateral end to its peak height. Black line: regression (linear model) across 1556 datasets with shaded background indicating 95% confidence interval. Pearson correlation coefficient per hemisphere: R = 0.64 (left hemispheric datasets; yellow) and R = 0.62 (right hemispheric datasets, red), p<.0001 indicates a strong correlation. Outer margins: Density (Gaussian kernel) of relative PPfpm height [%] (right), and relative PPfpm width [%] (upper) per hemisphere. **(B)** Hemispheric differences (left, right) in the relative PPfpm height [%] and relative PPfpm width [%]. Distribution of individual values is shown as a violin plot (Gaussian kernel) with .25 and .75 quartile indicated by horizontal lines; M ± SEM are given by hemisphere for dimensions of the PPfpm. Post-hoc paired t-test (Bonferroni correction: p_crit_<.0125) indicates a significant effect of hemisphere on dimension on PPfpm extent: **** p<.000025, *** p<.00025, ** p<.0025, * p<.0125, ns not significant. Yellow: left hemisphere; red: right hemisphere.

#### Left hemispheres exhibit a larger pli-de-passage fronto-pariétal moyen

The PPfpm - relative to the CS depth and length - tends to be higher than wider in both hemispheres (Table 10; Figure 9, B), though left hemispheres exhibit an overall higher and - to a lesser degree - also wider PPfpm (relative height: Δ = 3.41 %; relative width: Δ = 0.84 %; Bonferroni correction p_crit_ < .0125; Table 11).

**Table 10:** Mixed-effect ANOVA on extent (width/height) of the PPfpm (*n* = 1556 datasets).

| effect | $F(DF_{\text{num}}, DF_{\text{den}})$ | $p$ | |
| --- | --- | --- | --- |
| dimension | $F(1, 776) = 268.14$ | $< .0001$ | **** |
| hemisphere | $F(1, 776) = 38.70$ | $< .0001$ | **** |
| sex | $F(1, 776) = 5.25$ | .02 | * |
| dimension : hemisphere | $F(1, 776) = 42.33$ | $< .0001$ | **** |
| dimension: sex | $F(1, 776) = 3.02$ | .08 | ns |
| hemisphere : sex | $F(1, 776) = 0.81$ | .37 | ns |
| dimension : hemisphere : sex | $F(1, 776) = 1.25$ | .26 | ns |

Abbreviations: den, denominator; DF, degrees of freedom; num, numerator
Significance: \*\*\*\* $p < .0001$ , \*\*\* $p < .001$ , \*\* $p < .01$ , \* $p < .05$ , ns not significant

**Table 11:**
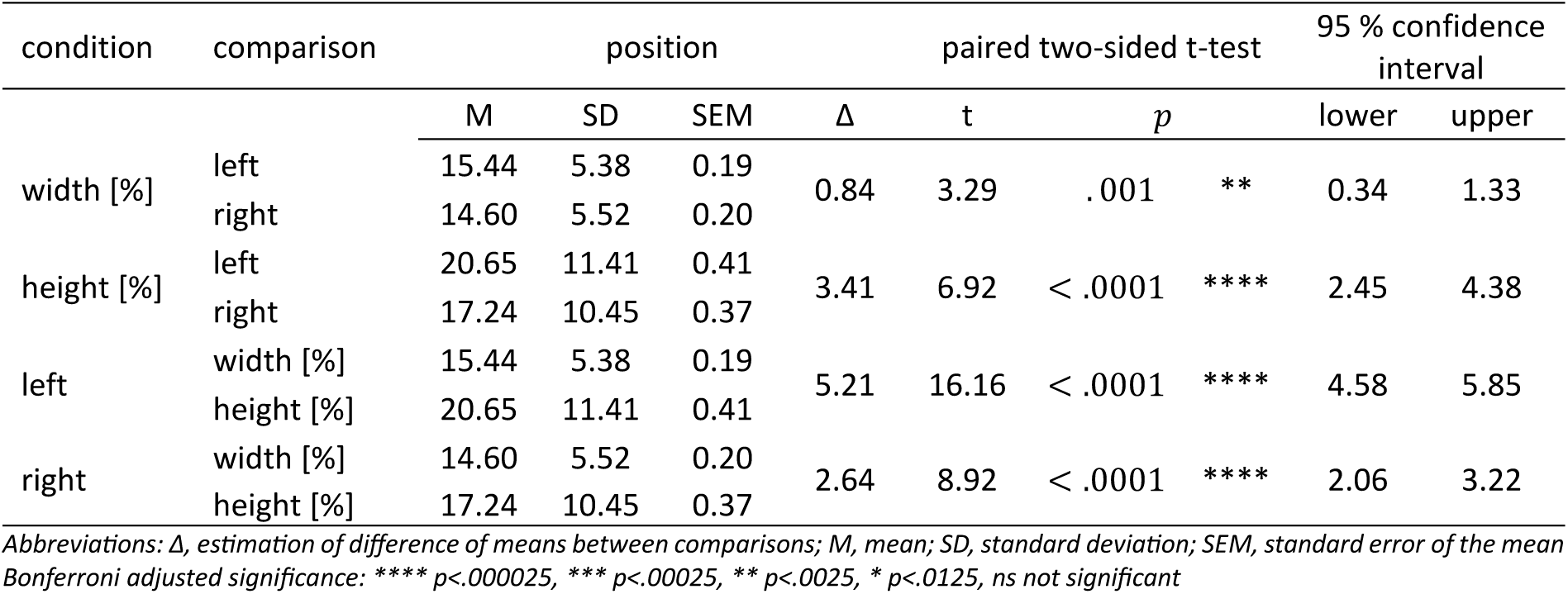
Bonferroni corrected post-hoc testing of PPfpm extent per hemisphere (*n* = 1556 datasets).

Together with the influence of hemispheres on PPfpm-I/-II, right hemispheres are indicated to - on average - exhibit flatter depth profiles correlating with a more lateral and smaller PPfpm with a lower peak height.

#### Relative height of the pli-de-passage fronto-pariétal moyen differs between males and females due to deeper sulci in males

For both PPfpm-I and PPfpm-II, depth but not positional values are significantly different between males (*n* = 374) and females (*n* = 404). Both PPfpm-I and PPfpm-II are deeper in males (PPfpm-I: Δ = 1.04 mm; PPfpm-II: Δ = 0.90 mm; Table 12), an effect suggested to originate from overall deeper CS in males (Figure 10, A).

**Figure 10:**
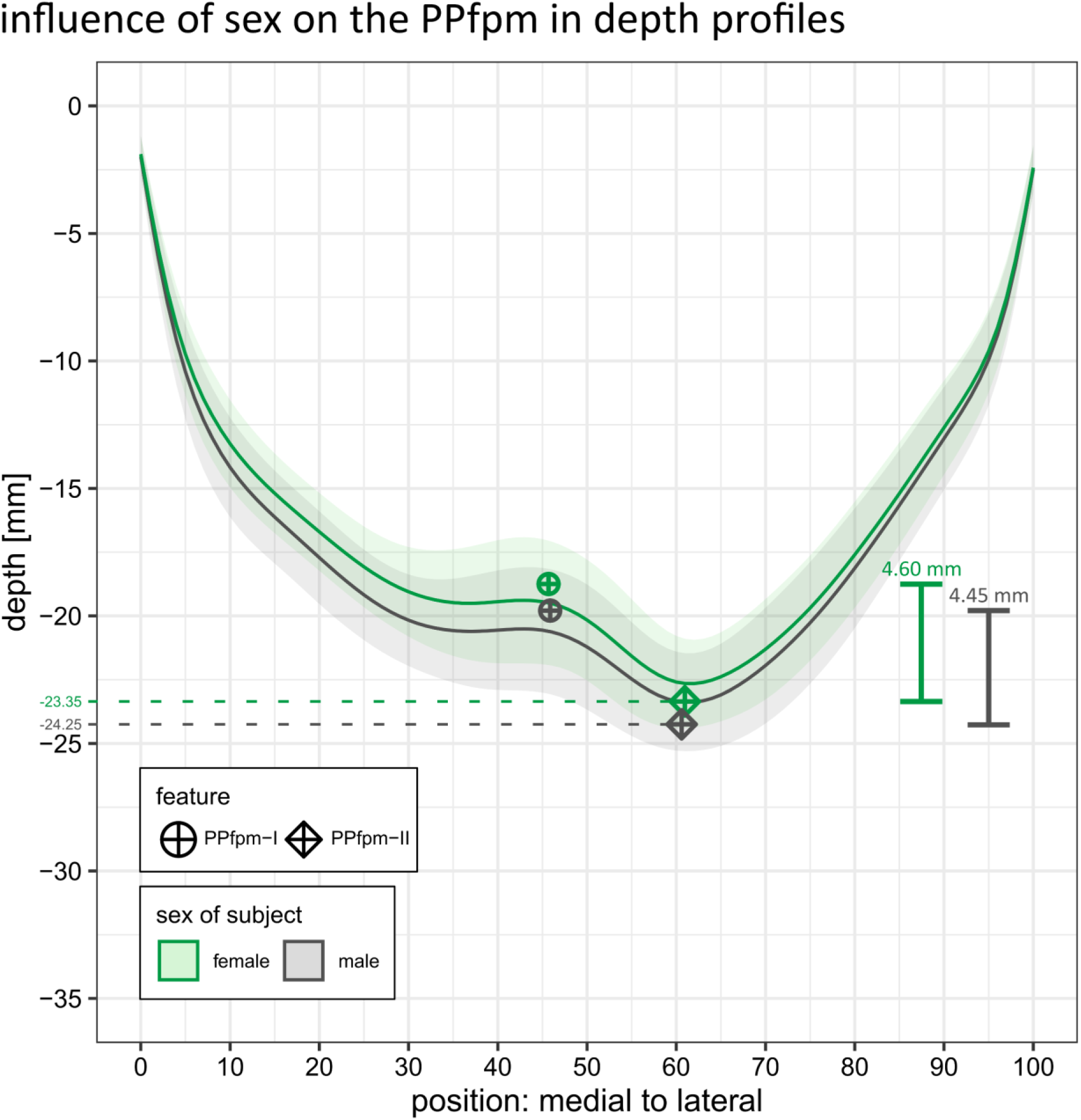
Effect of sex on the PPfpm in depth profiles. Average depth profile of males (grey, n = 748 datasets; n = 374 subjects) and females (green, n = 808 datasets; n = 404 subjects) computed per position; shaded area indicates the standard deviation. PPfpm-I (circle-cross) and PPfpm-II (diamond-cross) are averaged across all male and female datasets.

**Table 12:** Descriptive statistics of PPfpm extent and depth profile features PPfpm-I/-II by sex (*n* = 1556 datasets).

| sex |  | PPfpm extent |  |  | feature |  |  |  |
| --- | --- | --- | --- | --- | --- | --- | --- | --- |
|  |  | width [%] | height [mm] | height [%] | PPfpm-I |  | PPfpm-II |  |
|  |  |  |  |  | position | depth [mm] | position | depth [mm] |
| M | male | 14.74 | 4.45 | 18.23 | 45.87 | -19.79 | 60.61 | -24.25 |
|  | female | 15.28 | 4.60 | 19.60 | 45.69 | -18.75 | 60.96 | -23.35 |
| SD | male | 5.46 | 2.69 | 10.77 | 6.21 | 2.65 | 5.39 | 1.60 |
|  | female | 5.46 | 2.71 | 11.30 | 6.05 | 2.73 | 4.87 | 1.48 |
| SEM | male | 0.20 | 0.10 | 0.39 | 0.23 | 0.10 | 0.20 | 0.06 |
|  | female | 0.19 | 0.10 | 0.40 | 0.21 | 0.10 | 0.17 | 0.05 |

Abbreviations: M, mean; SD, standard deviation; SEM, standard error of the mean

A statistical effect of sex on the extent of the PPfpm is observed (Table 10). After supplementary two-way repeated measure ANOVA investigating the effect of hemisphere and sex on each the relative height and the relative width of the PPfpm, a week influence of sex is observed for relative PPfpm height only [“sex *F*(1, 776) = 5.03, *p* = .03], but not for relative PPfpm width [“sex”: *F*(1, 776) = 3.26, *p* = .07]. Two-way repeated measure ANOVA on the influence of hemisphere and sex on the absolute - in contrast to the relative - PPfpm height could not confirm any influence of sex [“sex”: *F*(1, 776) = 1.00, *p* = .32].

Given the observed depth difference in the depth trajectory between males and females and considering the effect of sex on the depth of PPfpm-I/PPfpm-II, the weak effect of sex on relative PPfpm height is implicated to result from the normalization procedure itself. That is, the absolute PPfpm height (Figure 10, B; Table 12) is not significantly different between males and females, while the relation to total CS depth at PPfpm-II is.

## Discussion

In the present study, structural MRI data from 1112 subjects of the HCP-YA S1200 Release were analyzed to provide a comprehensive macroanatomical characterization of the PPfpm in a large cohort: A method was devised to extract the PPfpm from CS depth profiles (*n* = 2165), defining two stable key features: the PPfpm’s lateral end (PPfpm-II: global minimum at position > 45) and its peak height (PPfpm-I: medially adjacent local maximum to PPfpm-II). The PPfpm was identified as a near universal cerebral fold at the CS fundus at mid-height: In depth profiles, both PPfpm-I and PPfpm-II are detectable in > 90 % of depth profiles (*n* = 1983); in structural MRI, the PPfpm was clearly detectable in all but three cases. Analyses confirmed the commonly small extent of the PPfpm < 5 mm; its height and width correlate positively. The PPfpm does, however, exhibit some variability in its shape and extent. Notably, the lateral end of the PPfpm coincides with the deepest point of the CS in > 95 % of depth profiles (*n* = 2079). A statistically significant, but minor, difference was found in the PPfpm’s position, on its peak height, and on its general extent between left and right hemispheres: On average, left hemispheres exhibit a more medial, higher, and slightly wider PPfpm. Subject sex had no effect on the absolute height of the PPfpm.

### Quality of extraction of the *pli-de-passage fronto-pariétal moyen* from central sulcus depth profiles

While the developed method accurately extracted the PPfpm from depth profiles in a majority of cases, there is a non-negligible number of datasets where extraction of the PPfpm was only partially possible (*n* = 182, 8.4 %) and/or incorrect (n = 146, 6.7 %).

Incomplete extraction was prevalent among very shallow depth profiles with a minuscule PPfpm (< 2 mm height) that did not allow for a precise identification of PPfpm-I. Additionally, the PPfpm of some depth profiles exhibited a steep incline towards medial, masked by the overall trajectory of the depth profile: Without a distinct peak height, PPfpm-I could thus not be identified by the implemented extraction method. Erroneous extraction of PPfpm-I or PPfpm-II resulted from only minor anatomical inaccuracies, wherein an overall flat depth profile impeded the identification of PPfpm-II as the deepest point, or where the deepest point of the CS did not match the lateral edge of the PPfpm. In some of these cases, refinement of the applied extraction method with a layered extraction based on a prior clustering of depth profile according to their main trajectory could have facilitated a more successful extraction. Refinement of the extraction method was, however, not pursued given that incomplete or erroneous extraction was prevalent in < 10 %, and given the robust quality control of the extraction method by one of the authors (AMM). To counteract a possible effect of erroneous/incomplete extraction on the detailed statistical analyses of the PPfpm, all analyses were performed on a subset of 1556 datasets, where a correct and complete extraction of the PPfpm was obtained in both hemispheres of a subject. Since characteristics of this subset (e.g. age of subjects, number of male/female subjects) showed no notable deviation from the complete dataset, statistical analyses are expected to be generalizable to the entire data under study.

### The shape of depth profiles and of the *pli-de-passage fronto-pariétal moyen* varies

As demonstrated, depth profiles exhibit considerable variation in their overall trajectories. This observation is consistent with the report by (McKay et al., 2013), who described three distinct depth profiles configurations: Unimodal depth profiles (prevalence: 8 %) lack a distinct central PPfpm, instead presenting a single deepest point accompanied by a sharp medial incline. This pattern closely resembles configurations observed in the current dataset, in which the PPfpm is embedded within a steep medial incline, occasionally complicating its identification, as discussed in detail above. Bimodal depth profiles are reported as most common (70 %) and display a clear, singular PPfpm elevation. This configuration corresponds to most cases observed in the present study, as exemplified in Figure 2 (B, upper left, lower left). Trimodal depth profiles (15 %) exhibited a double central elevation with multiples candidates for a discernible PPfpm elevation, a pattern found e.g. in Figure 2 (B, upper right, lower right). Although the categorization of depth profiles was not a primary objective of the current analysis, the observed data does not contradict the classifications proposed by McKay et al. (2013). However, the range of depth profile variability observed here suggests that such configurations may not represent discrete categories but rather points along a continuum that reflects the complex cortical topology of the human brain.

As implicated by McKay et al. (2013), and as shown in this study, the PPfpm itself presents most commonly as a small, singular elevation at central positions within depth profiles. However, a previously unreported portion of depth profiles and their PPfpm transition towards a double PPfpm elevation (< 10 %). Here, the PPfpm consists of not one singular, but of two more or less discrete peak heights separated by a local depression, seen e.g. in Figure 2 (B, middle column). Although these PPfpm shape variations may superficially resemble the “trimodal depth profiles” characterized by its two discrete elevations, careful examination of cortical reconstructions and the original structural MRI data reveals that they are, in fact, distinct with only one of these elevations corresponding to the PPfpm in depth profiles. The origin of this morphological variation remains unclear, and future studies should address this anatomical detail more thoroughly.

Additionally, this variability in PPfpm shape introduces ambiguity in the extraction of stable PPfpm features, particularly regarding the identification of the PPfpm’s peak height and its medial end, as multiple candidates may arise for both. To increase the consistency of the extraction of stable PPfpm features, the PPfpm peak height at PPfpm-I was thus defined as the maximum closest to the stable lateral end of the PPfpm at PPfpm-II. Extraction of the medial PPfpm end as a third feature was excluded from the analysis to mitigate potential confounding effects introduced by the highly variable double PPfpm and trimodal depth profile shape variants.

### The *pli-de-passage fronto-pariétal moyen* is a near-universal structure of commonly small extent located at mid-height in the central sulcus

The present study confirms that the PPfpm is a near-universal cerebral fold in the adult human brain. It was clearly detectable on either structural MRI and/or derived CS depth profiles in the vast majority of 1112 subjects.

Analyses of the PPfpm’s gross morphology highlights its generally small extent in the millimeter range with an average absolute height < 5mm and relative height < 19 %. Notably, a superficial PPfpm reaching the cortical surface and discontinuing the CS was found in 9 of 1112 subjects as reported separately by the authors (Schweizer et al., 2025). Results further demonstrated that the PPfpm height lies on a continuum, ranging from a subtle elevation to rare superficial variants. This aligns with the descriptions of Cunningham (1890), who described that the PPfpm can reach “all gradations between a mere shallowing […] of the [CS] and the presence of a distinct deep annectant gyrus”, and is consistent with a detailed report on the PPfpm height by Heschl (1877), replicated by the authors separately (Schweizer et al., 2025).

The average PPfpm width equals 15 % of total CS length. Due to the methodology employed for the CS parametrization, absolute CS length values cannot be calculated directly, as the measures are taken at discrete, relative positions along the CS’ medial to lateral trajectory. Thus, an exact metric measure of the PPfpm’s absolute width is unobtainable. However, with an average CS length of ca. 9.6 cm (Mashouf et al., 2017), the width of the PPfpm can be extrapolated to approximately 14.4 ± 5.2 mm, suggesting that the PPfpm is - in absolute measures - generally wider than higher. Notably, due to the lack of extraction of the PPfpm’s medial end, PPfpm width was defined as the relative distance from the PPfpm’s peak height to its lateral end, and not as the more intuitive distance between the PPfpm’s medial to lateral end. This methodological constraint should be considered when interpreting the findings.

Results further corroborate the consistent positional location of the PPfpm within the CS. Despite variability in extent and peak height, the PPfpm’s lateral end exhibited a remarkable positional consistency across this large cohort. On average, the PPfpm in depth profiles was consistently located at mid-height within the CS: the PPfpm’s peak height at PPfpm-I lies at position 46; the PPfpm’s lateral end at PPfpm-II at position 61. At first glance, this mid-height placement of the PPfpm appears to contrast with prior accounts that place the PPfpm at the superomedial third of the CS (Cunningham, 1890; Eberstaller, 1890; Skandalakis et al., 2025). However, this discrepancy may be attributable to methodological differences in defining and measuring the CS length. First, in the present study, depth measurements were taken at equidistant positions along the medial to lateral trajectory of the CS, thus compensating for individual variations in CS length and curvature. This level of standardization was likely difficult to achieve in earlier anatomical investigations, particularly those from the 19^th^ century. Second, the intrinsic variability of the CS curvature across individuals complicates visual estimations and may contribute to inconsistencies in previous reports on the PPfpm’s localization. Third, the medial and lateral extremities of the CS exhibit substantial anatomical variability, further impeding attempts to localize the PPfpm relative within the CS. In particular, the inferolateral end of the CS - near the lateral sulcus - has been reported to vary considerably between individuals (Eberstaller, 1890; Eichert et al., 2021), either being separated from the lateral sulcus by a large subcentral gyrus, or merging superficially with it due to confluence with the anterior subcentral sulcus, historically referred to as the “inferior transverse sulcus of the central sulcus” (“*Untere Querfurche der Centralfurche*”) (Cunningham, 1890; Eberstaller, 1890; Ecker, 1869; Turner, 1866). Taken together, while the present findings do not precisely replicate earlier descriptions of the PPfpm’s position within the CS, they are reconcilable with them when methodological and anatomical differences are considered.

### The *pli-de-passage fronto-pariétal moyen* in the context of the sulcal pits of the central sulcus

In the present study, extraction of the PPfpm from depth profiles was based on the identification of extrema - specifically a local maximum corresponding to the PPfpm’s peak height, and local/global minima corresponding to its medial and lateral end. Two prominent clusters of minima were thereby found to flank the average PPfpm in depth profiles at its medial and lateral ends (Figure 3). Although not an a priori goal of this study, these observations closely align with previous reports on the sulcal pits of the CS, prompting further consideration of the relationship between the PPfpm and sulcal pit anatomy.

Sulcal pits are defined as the locally deepest points of the cerebral cortex, located along the sulcal bottom lines within the sulcal basins of each cortical sulcus (Lohmann et al., 2008). They are proposed to be the first locations of cortical folding in the fetal brain, exhibit minimal change throughout cortical development (Lohmann et al., 2008; Régis et al., 2005), and are consistently described as stable anatomical landmarks in the human cerebral cortex. While the number of sulcal pits may vary across individuals and hemispheres, those within primary fissures such as the CS are reported to be stable in both number and position (Hostalet et al., 2025; Meng et al., 2014, 2018).

In the CS, two or three sulcal pits are typically observed in individuals, with the two-pit configuration being more common (Im et al., 2010; Meng et al., 2014, 2018). These two sulcal pits are usually located in the upper and middle third of the CS (Im et al., 2010; Meng et al., 2018), corresponding to the location of the PPfpm. Their position also aligns with anatomical descriptions by Cunningham (1890), confirmed by Régis et al. (2005) and Cachia et al. (2003), who describe the CS to arise from two primitive folds, or “sulcal roots”. It has therefore been proposed that these two most common CS sulcal pits correspond to the sulcal roots of the CS (Im et al., 2010; Meng et al., 2014). Considering that the PPfpm is described as the “eminence” between the CS sulcal roots, a close association between the PPfpm and the sulcal pits of the CS is both plausible and consistent with prior reports (Im et al., 2010; McKay et al., 2013). Thus, the medial cluster of minima is assumed to correspond to the first, upper sulcal pit, and the lateral cluster of minima is thought to correspond to the second, middle sulcal pit. In the present data, the lateral cluster of minima - from which landmark PPfpm-II was derived - was notably more stable in both position and depth, and coinciding with the deepest point of the CS. This finding is consistent with observations from Im et al. (2010), who reported a higher density of this sulcal pit. Notably, Cunningham (1890) described the lower sulcal root of the CS as forming earlier than the upper one, supporting the broader notion that sulcal pits are more stable the earlier they develop. This may further explain why the lateral sulcal pit - and thus the lateral cluster of minima - shows greater stability, whereas the medial cluster of minima was not sufficiently consistent to serve as a reliable reference for the medial end of the PPfpm in depth profiles.

Taken together, the present study reinforces the close anatomical association between the PPfpm and the CS sulcal pits. Notably, for other areas in the human brain, sulcal pits have further been closely linked to functional specialization (e.g., Auzias et al., 2015; Im et al., 2010; Lohmann et al., 2008; Natu et al., 2021; Régis et al., 2005). Given additional evidence on the association of the PPfpm to the sensorimotor hand/digit area (Alkadhi & Kollias, 2004; Boling & Olivier, 2004; Gordon et al., 2023; Jensen et al., 2023; Skandalakis et al., 2025), the need for further investigation on the structure-function relationship of the PPfpm, the CS sulcal pits, and functional regions in S1 and M1 is underscored.

### The *pli-de-passage fronto-pariétal moyen* displays stable features across hemisphere and sex

As outlined above, the PPfpm is a common and usually small structure in the human brain that exhibits some variation in extent and shape. While these variations are generally minor on a gross anatomical scale, the influence of biological factors, i.e., the hemispheric location of the PPfpm, and the sex of the subject, were analyzed:

Regarding the position of the PPfpm, the PPfpm’s peak height (PPfpm-I) and lateral end (PPfpm-II) were found overall more lateral in right hemispheres. While significant in this large data, the effect is minor with PPfpm-I more lateral by < 3 positions, and PPfpm-II more lateral by < 2 positions. Given reports on the different length of left (97.24 mm) and right hemispheric CS (94.85 mm) (Mashouf et al., 2017), this might reflect the different CS morphology of left versus right hemispheres rather than of the PPfpm itself. However, in the absence of the individual absolute length of the CS for this data, this hypothesis remains speculative. While statistically significant and highlighting intrinsic differences between left and right hemispheric CS, the observed effect lies, nevertheless, within millimeter range and thus underlines the PPfpm’s stability on a gross anatomical scale. Regarding depth values at both the PPfpm’s peak height and lateral end, the PPfpm’s peak height was found generally deeper in right hemispheres. While the depth at the PPfpm’s lateral end (PPfpm-II) remains consistent across all data and is unaffected by hemispheric location and the subject’s sex, the PPfpm’s peak height (PPfpm-I) was found to be significantly deeper in right hemispheres, although the difference was minimal at 0.82 mm on average. Together, this suggests that right hemispheres - on average - tend to exhibit a slightly smaller and - also given the correlation of PPfpm height with PPfpm width - narrower PPfpm that lies generally more lateral. Nevertheless, at the gross anatomical level, this effect is estimated to be < 1 mm and thus highlights the overall stability of the PPfpm.

Generally, the PPfpm was found to be unaffected in its absolute height by the subject’s sex with an average absolute height difference of 0.15 mm between males and females. In contrast, the relative PPfpm height was found significantly smaller in male brains, though the effect is minor (< 1.5 %). This difference is likely attributable to the CS being, on average, ca. 1 mm deeper at PPfpm-I/-II positions in males. Due to the absence of precise, modern data on the PPfpm during prenatal development and the lack of longitudinal information across the lifespan, it remains unclear when the PPfpm reaches its mature adult form, and whether this timing differs between sexes. While the consistent absolute height of the PPfpm across subjects is a noteworthy finding, its underlying origin remains uncertain.

In summary, while statistically significant, differences observed in the PPfpm’s extent, and on the position and depth at PPfpm-I/-II were all minor in magnitude. All reported effect sizes fall within, or only slightly exceed, the spatial resolution of the original MRI data, suggesting that these variations may, at least in part, reflect limitations in measurement precision. These findings thus reinforce that the PPfpm is a common, generally small, and structurally consistent cerebral fold within the human CS, and they highlight the remarkable macroanatomical stability of the PPfpm.

### Limitation

As the present study is conducted on a large dataset with > 1000 subjects from a diverse ethnic background, it is reasonable to assume that the presented results reflect the general characteristic of the PPfpm in the general population. Given that data were obtained exclusively from healthy subjects, the PPfpm’s structural characteristics might, however, differ in a patient population where organic brain disorders might affect the general brain morphology. Additionally, with the dataset restricted to young adults, it remains unknown how the PPfpm might be affected by age-related changes of cortical morphology. With ample evidence of a general decrease of cortical volume and surface in healthy aging adults, including the pre- and postcentral gyri (Fleischman et al., 2014; Zheng et al., 2019), it is reasonable to assume that the PPfpm itself might be affected by cortical thinning and decrease in total volume with increasing age. A translation of the PPfpm’s detailed characteristic to a clinical or aging population must thus be done cautiously. While the inclusion of additional structural MRI data, e.g. of the Chinese Human Connectome Project, the UK Biobank, or the Adolescent Brain Cognitive Development study, could have broadened the generalizability of the findings, the decision to focus exclusively on the HCP-YA data was made to ensure a high and consistent data quality and comparable preprocessing standards. Incorporating heterogeneous data would have introduced variability, potentially obscuring subtle structural features of the PPfpm. Further, a key strength of the present study is the rigorous quality control applied to each dataset individually, a standard that would be difficult to maintain in larger, more diverse populations.

A methodological limitation of the presented study, the detailed characterization of the PPfpm was conducted on computed CS depth profiles derived from structural MRI data and is not based on direct anatomical data. While this approach enables the precise and observer-independent characterization of the PPfpm in a large cohort (Cykowski et al., 2008), the resulting morphological and statistical analyses are derived from abstractions rather than direct anatomical observations. Minor differences between the MRI-based characterization of the PPfpm and those reported in historic (Heschl, 1877) and contemporary (Skandalakis et al., 2025) anatomical studies are thus likely. While a case-by-case validation of the PPfpm’s morphology in depth profiles versus true anatomical study would be ideal, it is - at present - unavailable. Nevertheless, the key findings of this study, i.e., the universality of the PPfpm, its small extent, its distinct shape as a cerebral fold, and its high positional stability across hemisphere and sex, are unlikely to be contradicted by a direct anatomical comparison. In fact, the general description of the PPfpm as a common structure with small but variable extent has recently been replicated across time in an independent sample (Schweizer et al., 2025).

### Anatomical location and identification of the *pli-de-passage fronto-pariétal moyen* in structural imaging data

To ease the identification of PPfpm on structural imaging data, this last section summarizes and describes how to best identify the PPfpm on structural MRI data on two exemplary subjects of the HCP-YA (for comparison of depth profile data see the same two subjects in Figure 5)

First, the PPfpm is generally located at mid-height in the CS and shows a close spatial relationship with the hand knob. On cortical surface reconstructions (Figure 11, left), it is located close to the middle frontal gyrus, and - depending on the overall trajectory and length of the CS - lies central or at the superomedial third of the CS. Generally, the PPfpm can be seen at the cortical surface only if of an unusually large extent (Figure 11, B), or in the rare case of a superficial PPfpm (Schweizer et al., 2025). Second, the PPfpm lies oblique and roughly perpendicular to the main trajectory of the CS (Heschl, 1877). With the CS following a main direction from superior to inferior, posterior to anterior, and medial to lateral, the PPfpm traverses from post- to precentral gyrus from inferior to superior, posterior to anterior, and lateral to medial. Given this oblique orientation of the PPfpm, only in some cases can it be observed in axial view, i.e., when of a large extent (Figure 11 B, middle) but not when of a small extent (Figure 11 A, middle), or when the CS presents with a less oblique main trajectory from superior to inferior. Third, the PPfpm can be best identified in sagittal view. Here it presents in the vast majority of cases as an unambiguous connection between pre- and postcentral gyrus traversing the CS: Thereby, the hook structure (Yousry, 1997) of the hand knob connects with a similar counter-hook on the postcentral side, forming a - in sagittal view - free-standing connection.

**Figure 11:**
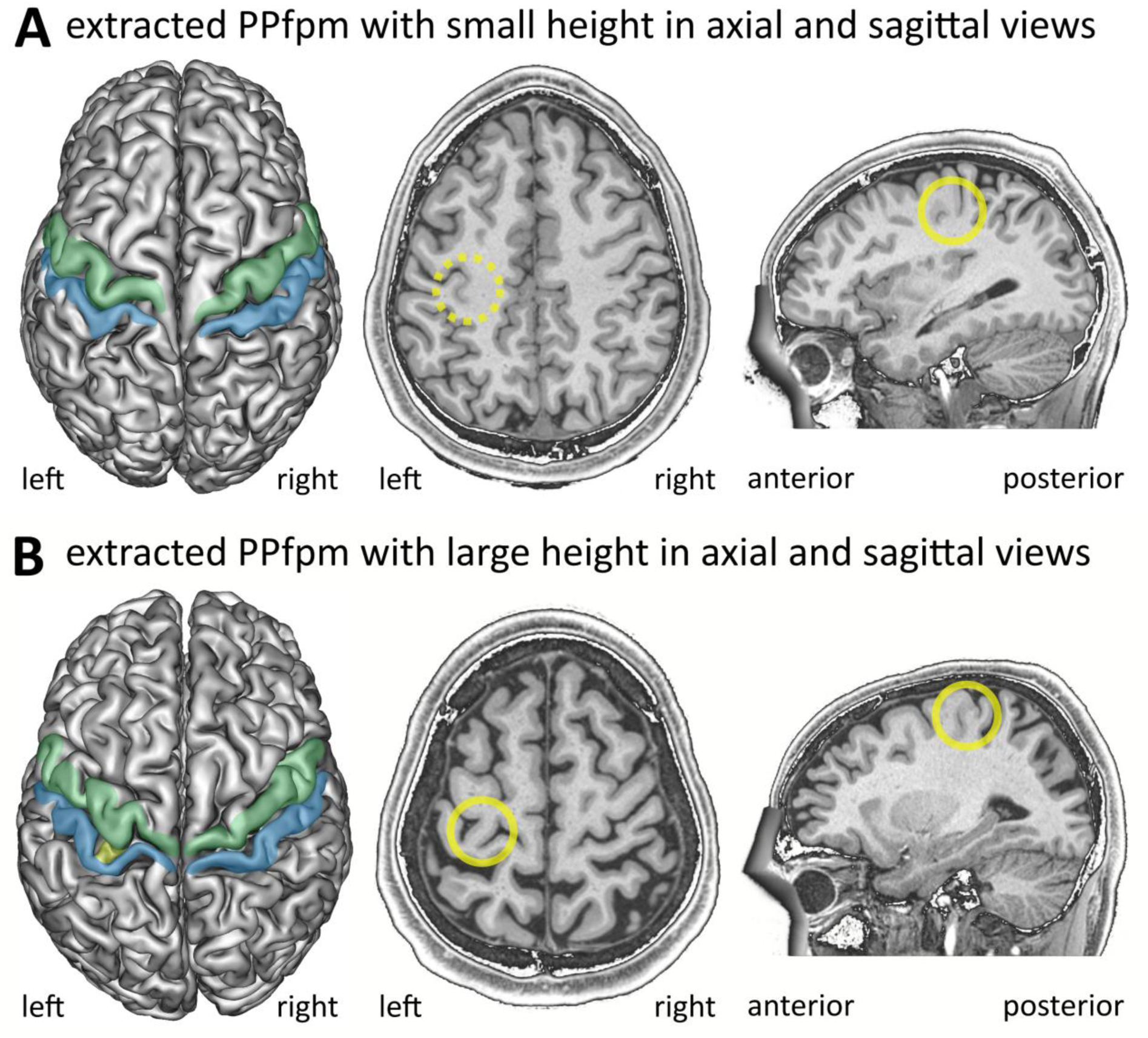
The PPfpm as seen on the cortical surface and in standard MRI views in native space. For two exemplary subjects of the HCP-YA, cortical grey matter reconstructions are shown in relation to the anatomy as seen in standard axial and sagittal MRI. Subjects display a PPfpm of **(A)** small and **(B)** large height in the left hemisphere. **Left:** Cortical grey matter reconstructions with marked precentral gyrus (green), postcentral gyrus (blue) and PPfpm (yellow). Note that the PPfpm is not visible in cortical grey matter reconstruction when its height is small but is apparent when its height is large (B). **Middle:** Axial MRI view with the PPfpm marked with a yellow circle. Note, that a PPfpm with a small height (A) is not clearly demarked in axial views; the dotted circle indicates the position of the PPfpm. Only a PPfpm with a large height (B) can be visualized as a distinct cerebral fold. **Right:** Sagittal MRI view with the PPfpm marked in a yellow circle. The PPfpm is generally detectable as a distinct connection between the precentral “hook” of the hand knob and a counter-hook of the postcentral gyrus. Note that the PPfpm can be detected in sagittal views even if it is of only a small height.

## Conclusion

In conclusion, the present study provides a comprehensive characterization of the *pli-de-passage fronto-pariétal moyen* (PPfpm), a deep cerebral fold hidden in the depth of the central sulcus (CS). Identified as a near-universal structure of the human brain, the PPfpm consistently exhibits two stable features: its peak height and its lateral end at the deepest point of the CS. Both features were subject to extensive quality control. By examining a large cohort of 1112 subjects of the Human Connectome Project Young Adult S1200 Release, the location, extent, and macroanatomical variability of the PPfpm are detailed. Demonstrating its remarkable consistency across individuals, hemispheres, and sex, the PPfpm emerges as a robust and easily identifiable structure in the human brain.

Together, this in-depth characterization of the PPfpm provides a solid anatomical framework for future research on the relationship of the PPfpm and the functional organization of the sensorimotor hand and digit area.

## Acknowledgements

Data were provided by the Human Connectome Project, WU-Minn Consortium (Principal Investigators: David Van Essen and Kamil Ugurbil; 1U54MH091657) funded by the 16 NIH Institutes and Centers that support the NIH Blueprint for Neuroscience Research; and by the McDonnell Center for Systems Neuroscience at Washington University. We acknowledge support by the Open Access Publication Funds of the University of Goettingen.

## Author contributions

AMM: conceptualization, data curation, formal analysis, funding acquisition, investigation, methodology, resources, software, validation, visualization, writing - original draft

RS: conceptualization, funding acquisition, project administration, resources, supervision, writing - review and editing

## Statements and Declarations

### Data availability statement

The structural magnetic resonance imaging data from the Human Connectome Project Young Adult S1200 Release that support the findings of this study are openly available at https://www.humanconnectome.org. Derived data are available from the corresponding author upon reasonable request.

### Ethics approval statement

Not applicable.

### Patient consent statement to participate

Not applicable.

### Permission to reproduce material from other sources

Not applicable.

### Clinical trial registration

Not applicable.

